# Observing the Cell in Its Native State: Imaging Subcellular Dynamics in Multicellular Organisms

**DOI:** 10.1101/243352

**Authors:** Tsung-Li Liu, Srigokul Upadhyayula, Daniel E. Milkie, Ved Singh, Kai Wang, Ian A. Swinburne, Kishore R. Mosaliganti, Zach M. Collins, Tom W. Hiscock, Jamien Shea, Abraham Q. Kohrman, Taylor N. Medwig, Daphne Dambournet, Ryan Forster, Brian Cunniff, Yuan Ruan, Hanako Yashiro, Steffen Scholpp, Elliot M. Meyerowitz, Dirk Hockemeyer, David G. Drubin, Benjamin L. Martin, David Q. Matus, Minoru Koyama, Sean G. Megason, Tom Kirchhausen, Eric Betzig

## Abstract

True physiological imaging of subcellular dynamics requires studying cells within their parent organisms, where all the environmental cues that drive gene expression, and hence the phenotypes we actually observe, are present. A complete understanding also requires volumetric imaging of the cell and its surroundings at high spatiotemporal resolution without inducing undue stress on either. We combined lattice light sheet microscopy with two-channel adaptive optics to achieve, across large multicellular volumes, noninvasive aberration-free imaging of subcellular processes, including endocytosis, organelle remodeling during mitosis, and the migration of axons, immune cells, and metastatic cancer cells *in vivo.* The technology reveals the phenotypic diversity within cells across different organisms and developmental stages, and may offer insights into how cells harness their intrinsic variability to adapt to different physiological environments.

**One Sentence Summary:** Combining lattice light sheet microscopy with adaptive optics enables high speed, high resolution *in vivo* 3D imaging of dynamic processes inside cells under physiological conditions within their parent organisms.

## Main Text

A common tenet, oft repeated in the field of bioimaging, is “seeing is believing”. But when can we believe what we see? The question becomes particularly relevant when imaging subcellular dynamics by fluorescence microscopy. After all, as powerful as genetically encoded fluorescent proteins have become, until recently they have rarely been used at endogenous expression levels, and therefore can upset the homeostatic balance of the cell. Furthermore, traditional imaging tools such as confocal microscopy are often too slow to study fast three-dimensional (3D) processes across cellular volumes, create out-of-focus photo-induced damage and fluorescence photobleaching, and subject the cell at the point of measurement (i.e., the excitation focus) to peak intensities orders of magnitude beyond that under which life evolved. Finally, much of what fluorescence microscopy has taught us about cellular processes has come from observing isolated adherent cells on glass, and yet it is certain that they did not evolve there. True physiological imaging requires studying cells within the organism in which they evolved, where all the environmental cues that regulate cell physiology are present.

Fortunately, a number of tools have been developed to address these problems. New genome editing technologies, such as CRISPR / Cas9 (*1*), enable expression of fluorescent fusion proteins at endogenous levels. Lattice light sheet microscopy (LLSM) (*2*) provides a non-invasive alternative for volumetric imaging of whole living cells at high spatiotemporal resolution, often over hundreds of time points. Adaptive optics (AO) (*3*) corrects for sample-induced aberrations caused by the inhomogeneous refractive index of biological specimens to recover resolution and signal-to-background ratio comparable to that seen for isolated cultured cells, even for cells deeply buried within multicellular organisms. The remaining challenge, then, is to combine these technologies in a way that retains their benefits and thereby enables the *in vivo* study of cell biology in conditions as close as possible to the native physiological state. Here we describe an adaptive optical lattice light sheet microscope (AO-LLSM) designed for this purpose, and demonstrate its utility through high speed, high resolution *in vivo* imaging of dynamic subcellular processes in 3D, including clathrin-mediated endocytosis, organelle remodeling during mitosis, and the migration of axons, immune cells, and metastatic cancer cells in crowded multicellular environments.

## Lattice Light Sheet Microscope with Two-Channel Adaptive Optics

Although several approaches to AO have been demonstrated in biological systems (*3*), including in the excitation (*4*) or detection (*5*) light paths of a light sheet microscope, we chose an approach where the sample-induced aberrations affecting the image of a localized reference “guide star” created via two-photon excited fluorescence (TPEF) within the specimen are measured and then corrected with a phase modulation element (*6*). By scanning the guide star over the region to be imaged (*7*), an average correction is measured that is often more accurate than single point correction – essential, since a poor AO correction is often worse than none at all. Scanning also greatly reduces the photon load demanded from any single point. Coupled with correction times as short as 70 ms (*7*), these advantages make this AO method compatible with the speed and noninvasiveness of LLSM.

In LLSM, light traverses different regions of the specimen for excitation and detection, and therefore is subject to different aberrations. Hence, independent AO systems are needed for each. This led us to design a system (Fig. 1A, supplementary note 1, fig. S1) where infra-red light (red) from a Ti:Sapphire ultrafast pulsed laser (2PL) is ported to either the excitation or detection arm of a LLSM (left inset, Fig. 1A) by a switching galvanometer (SG1). In the detection case, TPEF (green) generated within a specimen by scanning the guide star across the focal plane of the detection objective (DO) is descanned (*7*) and sent to a Shack-Hartmann wavefront sensor (DSH) via a second switching galvanometer (SG2). We then apply the inverse of the measured aberration to a deformable mirror (DM) placed conjugate to both DSH and the rear pupil plane of DO (supplementary note 2). Because the signal (also green) generated by the lattice light sheet when in imaging mode travels the same path through the specimen as the guide star, and reflects from the same DM, the corrective pattern we apply to DM produces an AO corrected image of the current excitation plane within the specimen on the camera (CAM) when ported there by SG2.

**Fig. 1.**
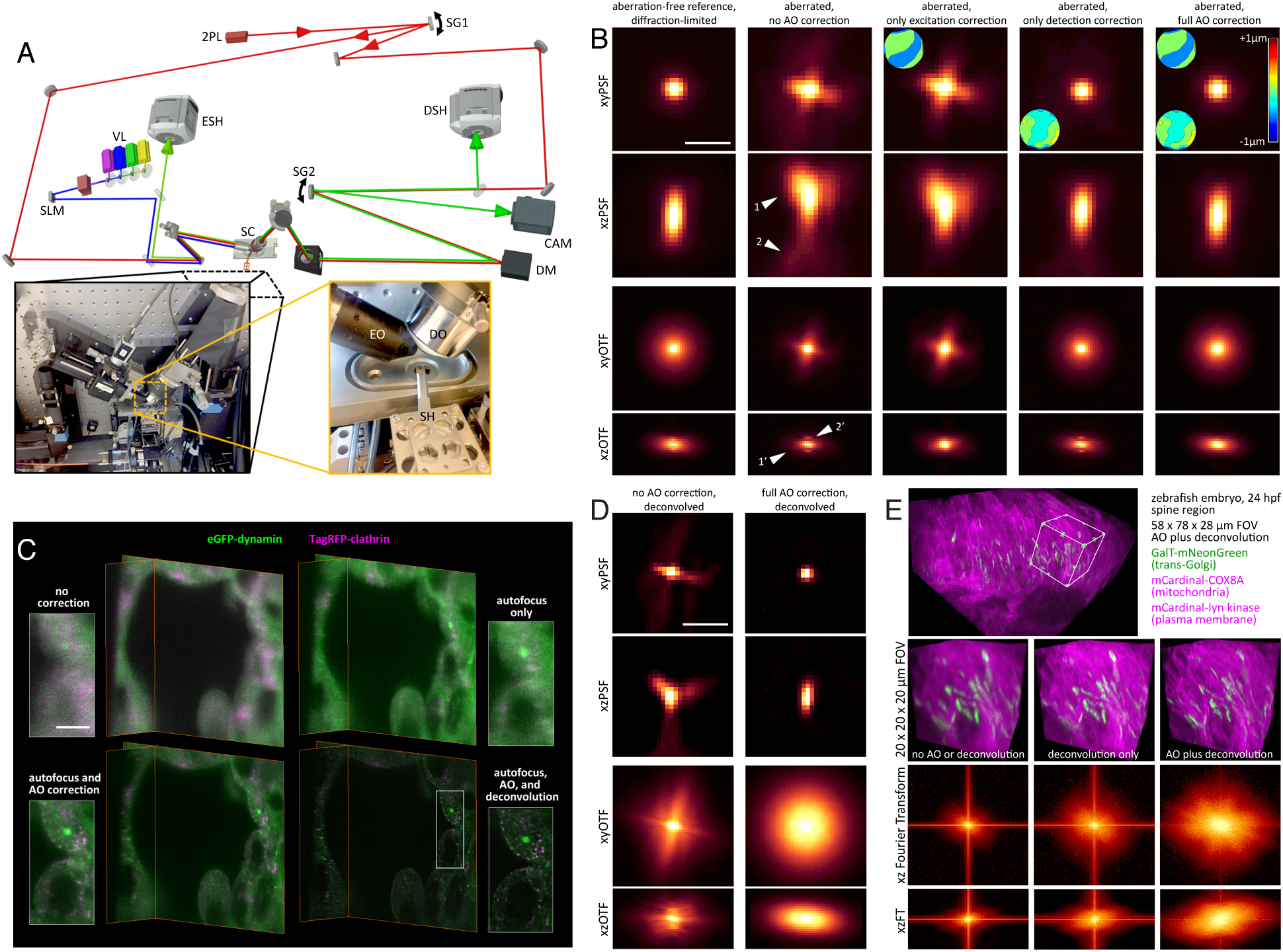
Adaptive Optical Lattice Light Sheet Microscope (AO-LLSM). (**A**) Simplified schematic of the microscope (*c.f*, fig. S1). (**B**) Orthogonal xy and xz maximum intensity projections (MIPs) of the point spread function (PSF, top two rows) and corresponding optical transfer function (OTF, bottom two rows) of the microscope under five different degrees of AO correction (columns), as measured from a 100 nm diameter fluorescent bead in an aberrating agarose gel. Insets show the corrective wavefronts applied, where appropriate. Scale bar, 1 μm. (**C**) Orthogonal orthoslices for four different levels of correction as shown within a live human stem cell-derived organoid grown in Matrigel, gene-edited to express endogenous levels of clathrin and dynamin in coated endocytic pits (*c.f.*, Movie 1). Scale bar, 5 μm. (D) MIPs and corresponding OTFs of the uncorrected (left column) and fully corrected (right) bead images from Fig. 1B, after deconvolution using the aberration-free reference PSF. Scale bar, 1μm. (E) Cellular trans-Golgi, mitochondria, and plasma membranes in the spine of a live zebrafish embryo 24 hpf, showing unprocessed data without AO correction (left column), deconvolved data without AO correction (center), and deconvolved data after AO correction (right) (*c.f.*, movie S2). Orthogonal MIP views (bottom two rows) of the Fourier Transform (FT) of the data in all three cases indicate the degree of information recovery after AO correction with deconvolution.

Similarly, for excitation correction, we send descanned TPEF generated and collected through the excitation objective (EO) to a second wavefront sensor (ESH). However, because LLS excitation is confined to only a thin annulus at the rear pupil of DO (*2*), a DM placed conjugate to this pupil would be ineffective for AO correction over most of its surface. Instead, we apply wavefront correction at the same sample-conjugate spatial light modulator (SLM) which creates the light sheet itself, thereby enlisting thousands of independently corrective pixels. To do so, we subtract the measured phase aberration from the phase of the Fourier transform (FT) of the ideal, aberration-free SLM pattern, and then inverse transform the result back to the sample-conjugate SLM plane (supplementary note 3, fig. S2).

Finally, optimal resolution requires the LLS to be coincident with the focal plane of DO to less than ~0.46 μm over the entire field of view (FOV), whereas refractive index differences between the specimen and the surrounding media lead to tip-tilt alignment or axial displacement errors for the light sheet that might exceed this. Fortunately, we find both empirically and by estimation (supplementary note 4, fig. S3) that only displacement is a concern over FOVs typical of LLSM, and can be robustly corrected by imaging, edge-on through EO, the plane of TPEF we generate when measuring the detection aberration, and then measuring its offset relative to plane of fluorescence generated by the LLS, also viewed edge-on through EO (supplementary note 5, fig. S4).

As an example (Fig. 1B), after correction for aberrations in the microscope itself, the point spread function (PSF) and corresponding optical transfer function (OTF), measured from a 200 nm bead, indicate nearly diffraction-limited performance (column 1). However, when a similar bead is placed in 2% low-melt agarose, we observe substantial aberration (column 2), both laterally (arrows 1, 1') and axially (arrows 2, 2'). Correcting only the excitation aberration improves the axial resolution (column 3) by returning the light sheet to its original width (fig. S2). Conversely, correcting only the detection aberration improves primarily the lateral performance (column 4). However, when we combine excitation and detection correction, the image of the bead is returned to its diffraction-limited form (column 5), with an eight-fold recovery to its original peak intensity. Furthermore, the same correction proves valid over a 30 μm field of beads in agarose (movie S1).

Next, we imaged organoids differentiated from human stem cells, gene edited to express endogenous levels of TagRFP-clathrin and eGFP-dynamin in clathrin coated pits (CCPs). Such organoids permit the study of human cellular differentiation and tissue morphogenesis at subcellular resolution, with an experimental accessibility difficult to achieve *in vivo.* However, the extracellular matrix in which they are grown introduces significant aberrations, while the fast dynamics and limited number of fluorophores in each CCP demand high speed imaging with low photobleaching. The system is therefore well suited to demonstrate the capabilities of AO-LLSM for live multicellular imaging, with the CCPs doubling as broadly distributed puncta of sub-diffractive size to evaluate imaging performance.

Without any AO or focus correction (upper left of Fig. 1C and Movie 1), no CCPs are visible, and the cell boundaries are poorly defined. Autofocus alone (upper right) reveals larger clusters of CCPs, but must be combined with complete AO correction (excitation and detection, lower left) to identify individual CCPs. At that point, the imaging is diffraction limited, so the data can be deconvolved using the system corrected PSF to compensate for the spatial filtering properties of the microscope. Both dynamin and clathrin puncta then stand out clearly above the cytosolic background (lower right), allowing us to quantitatively measure their lifetimes and diffusion tracks in 3D for 120 time points at 1.86 sec intervals (Movie 1). The increasing recovery of information as we progress through these four cases can also be seen quantitatively in their corresponding OTFs (fig. S5).

One of the key advantages of complete AO correction is that it enables accurate deconvolution (Fig. 1D), giving the most truthful representation of the specimen possible within the limits of diffraction (column 2). In contrast, applying deconvolution to an aberrated image gives a distorted result, because the OTF can then fall below the noise floor asymmetrically and at spatial frequencies well below the diffraction limit (column 1), after which they cannot be recovered by deconvolution. These same trends can be seen in densely labeled specimens as well, such as across mitochondria, Golgi, and PM of cells near the spinal midline in a living zebrafish embryo 24 hours post fertilization (hpf) (Fig. 1D, right half and movie S2), where the greatest information content as seen in the FT of image volume is obtained by combined AO and deconvolution.

## Dynamics of Clathrin-Mediated Endocytosis in Zebrafish

Whereas for multicellular studies of human cells we are usually confined to *in vitro* systems such as organoids, for transparent model organisms we can apply AO-LLSM *in vivo*, where the complete physiological environment is preserved. In particular, the zebrafish has become the most widely used non-mammalian model vertebrate organism. We took advantage of its small size and transparency to visualize, in real-time, the formation of endocytic clathrin-coated pits and clathrin-coated vesicles (CCVs) in the context of the developing organism. We first imaged a 74.5 × 99.3 × 40.6 μm volume of developing muscle near the dorsal side of the tail in a fish larva at 80 hpf stably expressing clathrin light chain A fused to monomeric dsRed (*8*) at 10 sec intervals for 15 min (Fig. 2A, Movie 2). We observed a high spatial density of diffraction-limited clathrin-spots that, after computational separation of all cells, were mostly colocalized with the plasma membrane (PM). As expected, these spots appeared and disappeared with the formation of new CCPs and their eventual uncoating. We determined that the areal density of CCPs was similar in muscle fibers (e.g., green arrows, Fig 2A) and the endothelial cells (magenta arrows) lining blood vessels (0.13 vs 0.17 CCPs/μm^2^, respectively) but the distribution of their intensities, as measured before the onset of noticeable dsRed photobleaching, was broader in the endothelial cells (13 vs. 36 counts median absolute deviation (MAD), lower inset, Fig. 2A), with the brightest endothelial pits skewing the median intensity higher (30 vs. 49 counts). Since CCP size is proportional to intensity (*9*), these results indicate a broader range of CCP sizes in the vasculature endothelium, including large structures, which may include unresolved clusters of CCPs (*10*), up to six-fold larger than the median size in muscle fibers.

**Fig. 2.**
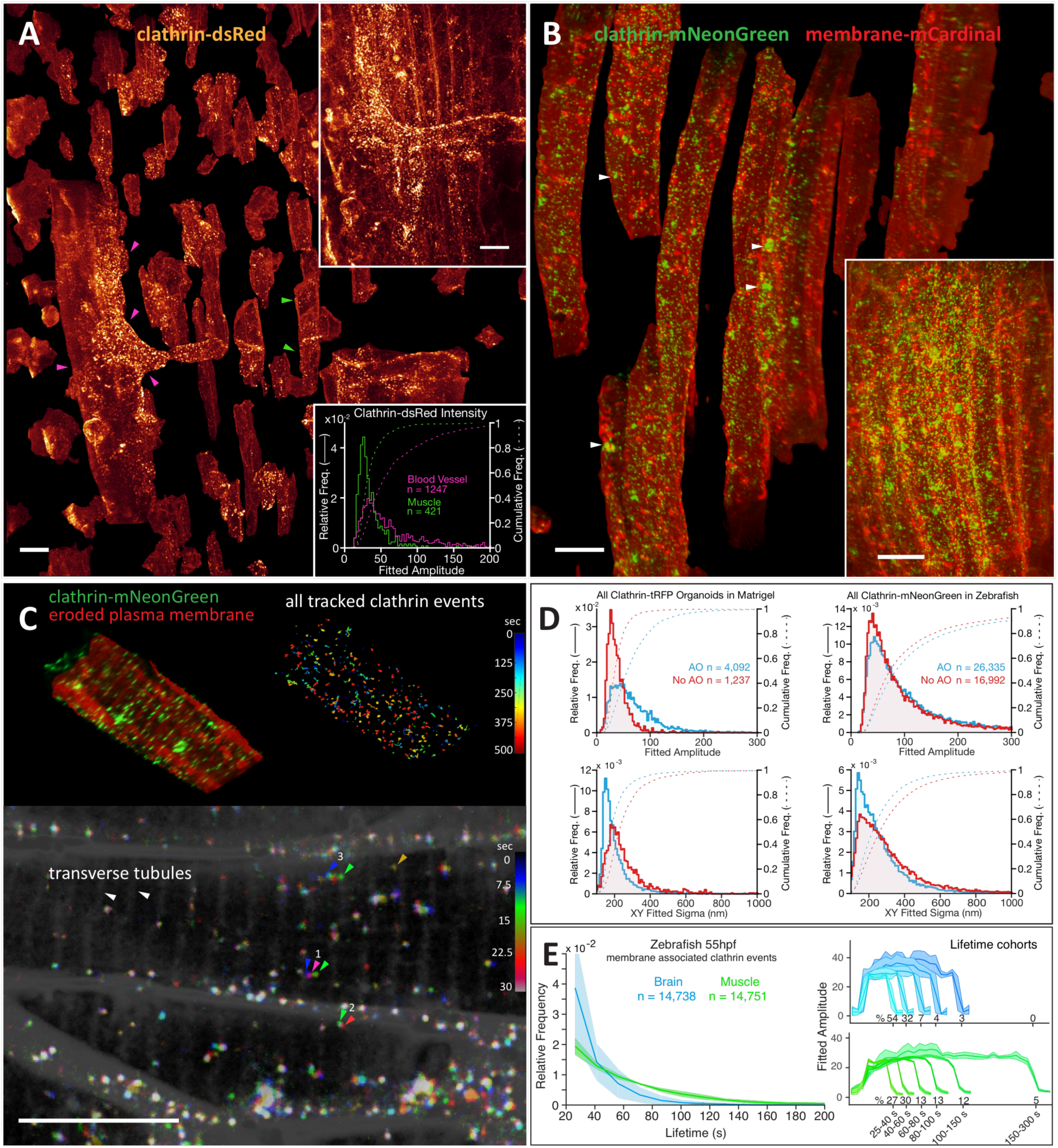
Clathrin-mediated endocytosis in zebrafish. (**A**) Volume rendering of computationally separated muscle fibers (e.g., green arrows) and vascular endothelial cells (e.g., magenta arrows), both expressing clathrin-LCA-DsRed to highlight clathrin-coated pits (CCPs), and vesicles (CCVs) from muscle tissue (upper inset) in the dorsal tail region of a developing zebrafish larva 80 hpf (*c.f*, Movie 2). Brighter clathrin puncta were observed in the endothelial cells (lower inset). Scale bars, 10 μm. (**B**) Computationally separated muscle fibers from a region (lower inset) in the tail of a zebrafish embryo 50 hpf, displaying both individual CCPs and CCVs as well as larger clathrin-rich vesicles (arrows) (*c.f.*, Movie 3). Scale bars, 10 μm. (**C**) Spatial distribution and dynamics of CCPs and CCVs tracked for 12 minutes at 7.5 sec intervals in one muscle fiber from Fig. 2B, showing CCPs localized at t-tubules (top left) as well as diffusion and lifetime characteristics for CCPs and CCVs across the cell (top right). A MIP through a 2 μm thick slab and three consecutive time points (bottom) shows examples of a pinned CCV (arrows 2) and slowly diffusing (arrows 1) or rapidly shuttling CCVs (arrows 3) (*c.f*, movie S3). Scale bar, 10 μm. (**D**) Effect of AO on the measured quantity, intensity, and localization precision of CCPs and CCVs in the organoid in Fig. 1C and the zebrafish in Fig. 2B. (**E**) Comparative distribution of CCP / CCV lifetimes in the brain and muscle of a developing zebrafish embryo 55 hpf.

To track CCPs over their entire lifetimes without excessive photobleaching, we then turned to zebrafish embryos expressing, via mRNA injection, the brighter and more photostable fluorescent protein mNeonGreen fused to clathrin light chain A. These embryos also ubiquitously expressed the lyn kinase membrane-targeting motif fused to mCardinal, facilitating the computational separation of all cells throughout the organism. Embryos imaged 50–55 hpf in muscle fibers at the tail (Fig. 2B, fig. S6A, Movie 3) and in the hindbrain (fig. S6B) displayed a variety of morphologies, trafficking behaviors, and lifetime distributions. Large (micron-scale and above) intracellular spots of limited mobility (arrows, Fig. 2B) probably represent clathrin-rich vesicles clustered at the trans-Golgi network, while smaller and more mobile internal diffraction limited objects (Fig. 2C, arrow group 1) likely correspond to endosomal carriers similar to those seen in cultured mammalian cells (*10*). On the other hand, diffraction limited spots we observed at the PM (Fig. 2C, arrow group 2) likely represent individual CCPs and CCVs. We also found apparent CCPs at the t-tubules spanning muscle fibers (Fig. 2C, top left), in contrast to the diffuse clathrin signal seen via immunofluorescence in fixed rat muscle fibers (*11*). Most of these were pinned at one location, but on occasion they would break free from a t-tubule and exhibit fast displacement along the fiber axis (Fig. 2C, arrow group 3, and movie S3), possibly indicative of active transport of CCVs along myofibrils or within the sarcoplasmic reticulum.

The increased brightness and photostability afforded by mNeonGreen meant that we could find a much greater proportion of bright CCPs that we could track for their entire lifetimes (Fig. 2D). AO then further allowed us to detect more CCPs (fig. S7) and track all CCPs with higher precision (Fig. 2C, top right, fig. S8A-C, and Movie 3). Comparing CCPs in muscle fibers and the brain, we found that, although their initiation frequencies and fluorescence intensities were similar, brain endocytic-coated pits and vesicles formed relatively faster (Fig. 2E). Assuming that clathrin puncta in muscle and brain cells lasting at least 21 sec corresponded to successful coated vesicles, each with an assumed membrane diameter of 60 nm, we estimate that ~0.1% of the PM is internalized through the clathrin pathway every minute (fig. S8D). This is similar to the values derived from initiation rates of bona fide coated pits in cultured SUM-159 (*12*) or htertRPE-1 (*13*) mammalian cells at 37°C.

## *In Vivo* Imaging of Organelle Morphology and Dynamics

A major focus in cell biology is the study of the structure and function of organelles within the living cell. To study three organelles simultaneously across a population of cells *in vivo*, we injected mRNAs into a zebrafish embryo at the 1–2 cell stage to express mNeonGreen-GalT and tagRFP-Sec61β as markers of the trans-Golgi and endoplasmic reticulum (ER), respectively. We then additionally stained the embryo with the live cell dye MitoTracker Deep Red to visualize mitochondria, and chose an embryo line that transgenically expressed membrane-targeted Citrine to aid in the computational segmentation and separation of all cells, so that we could study them individually (Fig. 3A).

**Fig. 3.**
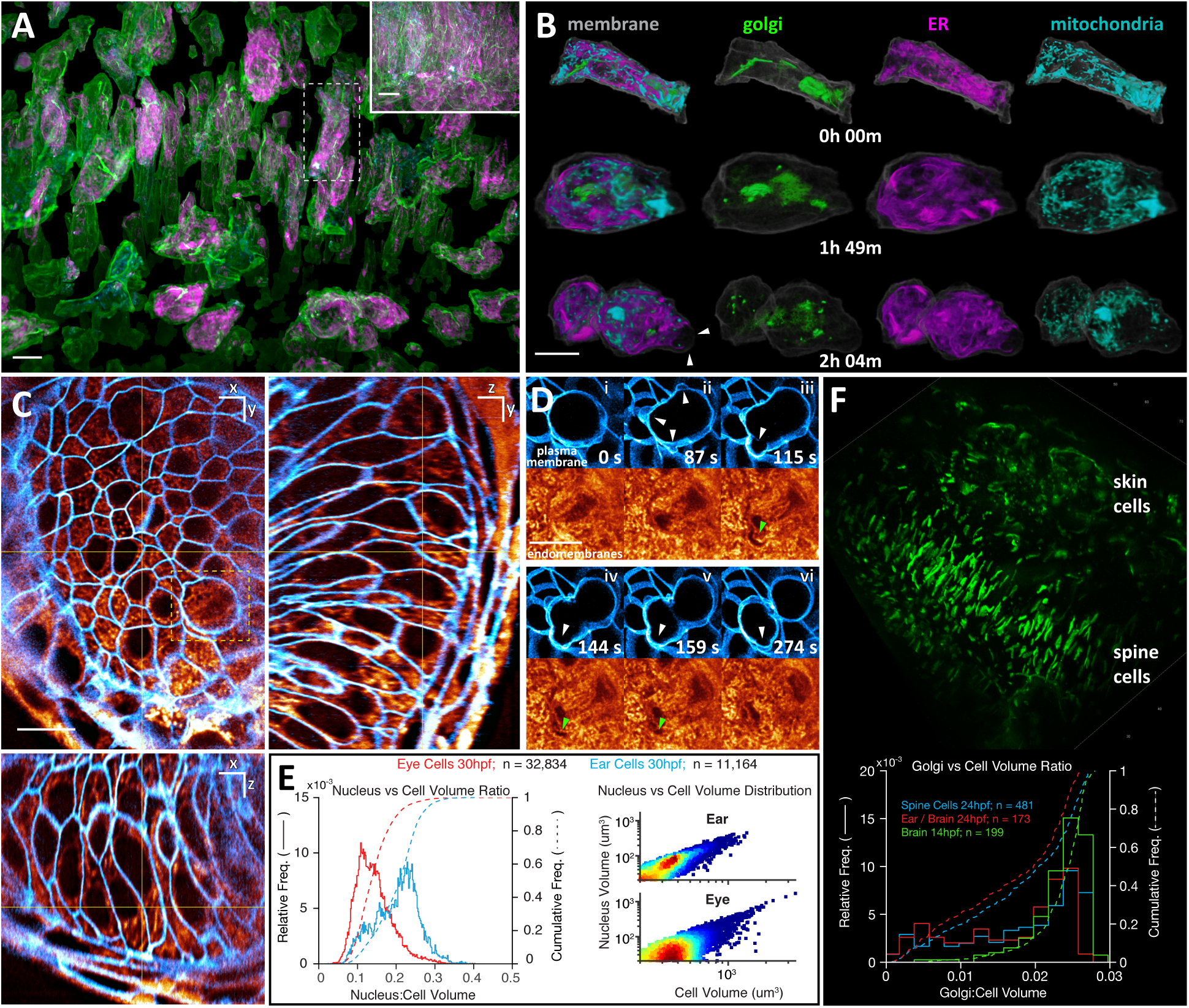
Organelle morphologies and dynamics in zebrafish. **(A)** Computationally separated neural progenitor cells from a region (inset) in the developing brain of a zebrafish embryo imaged from 14.0 to 16.5 hpf at 44 sec intervals (*c.f*, Movie 4). Scale bars, 10 μm. **(B)** Changing morphologies of three organelles in the cell outlined in Fig. 3A, from interphase (top row) through metaphase (middle row) to cytokinesis (bottom row). **(C)** MIP views from 1 μm thick orthogonal slabs within the eye of a zebrafish embryo 30 hpf, showing fluorescently labeled plasma membranes (blue) and endomembranes (orange) (*c.f*, Movie 5). Scale bars, 10 μm. **(D)** Six time points from Movie 5, showing membrane blebs (white arrows) during mitosis, and the exclusion of endomembranes in blebs (green arrows). **(E)** Relationships between nuclear volume and total cell volume in the eye and ear, 30 hpf. **(F)** Different morphologies of trans-Golgi (top) near the spine of a zebrafish embryo 24 hpf, and distribution of trans-Golgi volume in different cell types and at different developmental stages.

Imaging brain progenitor cells from 14 hpf across a 70 × 35 × 35 μm volume for 2.5 hours at 44 sec intervals allowed us to follow organelle morphology throughout the cell cycle (Movie 4). In interphase, we observed multiple trans-Golgi segments in most cells, often appearing as long filaments preferentially aligned along the axis of cell polarization (Fig. 3A) that disassembled into a diffuse haze or small fragments during mitosis (Fig. 3B). The ER largely recapitulated its common form in cultured cells: a reticular network in interphase, and sheet-like cisternae during mitosis (*14*). Dye uptake by mitochondria was rather sparse, but revealed largely punctate structures near the surface, and longer tubules, familiar from cultured cells, in the subset of internal interphase cells that were well-labeled. Analyzing one such cell, we found that the distribution of all three organelles was fairly uniform from the PM to the nucleus in interphase (fig. S9A), but the mitochondria were preferentially located nearer the PM during mitosis (fig. S9A, 109 min).

The early synchrony of cell division is lost in zebrafish at the midblastula transition (3 hpf). Nevertheless, at 14 hpf we observed instances of cascading cell division, where adjacent cells underwent mitosis one after another (fig. S10 Movie 4). Mitotic cells, as observed previously in cultured cells (*12*), decreased their surface area (fig. S9B) as they assumed a spheroidal shape prior to cytokinesis, but then recovered their initial area after division. Total cellular volume remained constant throughout mitosis (fig. S9B). Notably, however, we observed instances of asymmetric cytokinesis (Fig. 3B, Movie 4), where the two daughter cells had different shapes and surface areas. We also found, consistent with isolated cultured cells (*12*), that the rounded mitotic cells seen here (Movie 4) and in the developing eye 30 hpf (white arrows in Fig.3D, Movie 5) produced transient blebs prior to cytokinesis. However, unlike cultured cells, these cells exist in a dense multicellular environment where blebs must generate substantial force to displace adjacent cells. Indeed, by labeling all intracellular membranes in the eye specimen with Bodipy-TMR, we found that the hydrostatic pressure necessary to creates the blebs initially excluded the endomembranes, which then only slowly filled the newly created void (green arrows in Fig. 3D).

Lastly, we observed considerable variability in the size and morphology of specific subcellular features across different organs and developmental stages. Bodipy-TMR negative staining (e.g., Fig. 3C, D) revealed that the nuclei of ear cells 30 hpf were nearly twice as large as those of the eye when normalized by the total cellular volume (Fig. 3E, left), and that nuclear volume and cellular volume were positively correlated (Fig. 3E, right), with Pearson correlation coefficients of 0.90 for the ear and 0.80 for the eye. Likewise, we found that the median normalized trans-Golgi volume in brain progenitor cells 14 hpf was significantly larger (2.46%, MAD = 0.26%) than in ear, brain, or spine cells (2.0%, MAD = 0.66%) 24 hpf, although in all cases the trans-Golgi comprised less than 3% of the total cellular volume (Fig. 3F). Golgi took many forms, from the aforementioned narrow polarized filaments in the early brain (Fig. 3A), to shorter segments clustered near the midline in the spine and nuclear-wrapping filaments in skin cells (Fig. 3F).

## Tiled Acquisition for Aberrations Varying in Space and Time

Because the refractive index profile can vary across a specimen and can also vary as the specimen develops, the AO corrections required can vary in both space and time. Unfortunately, it is difficult estimate *a priori* the size or temporal stability of the isoplanatic patch – the FOV over which a single AO correction is valid. Empirically, we have found in zebrafish embryos less than 72 hpf that a single excitation/detection correction pair obtained by scan/descan over the FOV is usually valid across 30–60 μm in each direction for at least 1 hr, provided that the light does not intersect the yolk. Fortuitously, these dimensions are comparable to those over which a LLS of sub-micron thickness does not deviate substantially in width. The examples shown above largely fall within these limits, and hence for them a single AO correction pair at a single time point sufficed.

In other cases, however, we may wish to cover much larger FOVs, such as to study variability across a large population of cells, collective cell migration, or organogenesis. To do so, we must stitch together data from multiple image subvolumes, each with its own independent AO correction. To demonstrate the necessity of this, we imaged a 213 × 213 × 113 μm volume (Fig. 4A and movie S4) comprised of 7 × 7 × 3 subvolumes in the tail region of a zebrafish embryo 96 hpf by three different protocols: no AO correction (Fig. 4B, column 1); AO correction from the center tile applied to all tiles (column 2); and independent AO correction in each tile (column 3). When viewed across 3 μm thick slabs perpendicular (*xy*, Fig. 4B, top row) or parallel (xz, bottom row) to the detection axis, a small volume within the center tile (small orange boxes) showed substantial improvement by either center tile or all tiles AO correction, in both the xy lateral (upper left, top row) and xz axial (upper left, bottom row) planes. This is to be expected, since the site of AO correction coincides with the viewing area in these two cases. However, in a small volume at the edge of the fish (small blue boxes), only the data taken using individual AO corrections in each tile recovers optimal resolution in all directions (lower right blue boxes, column 3), because this volume is outside the isoplanatic patch over which the center tile AO correction is valid. Indeed, applying the center correction across the larger stitched volume often results in greater wavefront errors (fig. S11) and poorer resolution (lower right blue boxes, column 2) than applying no correction at all (lower right blue boxes, column 3), highlighting the importance of accurate and robust correction, if AO is to be applied at all.

**Fig. 4.**
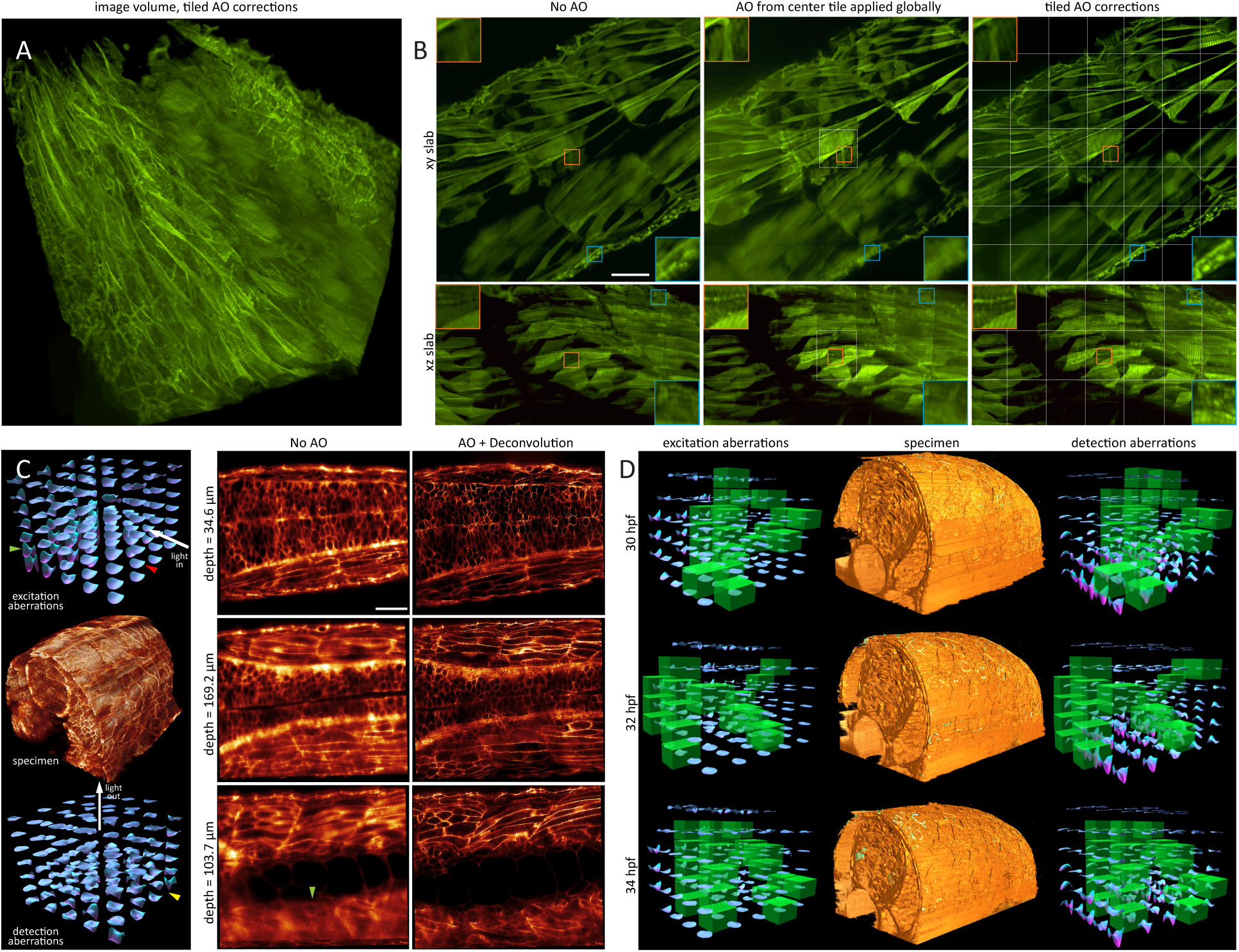
AO-LLSM over large volumes. (**A**) Aberration corrected volume rendering over 213 × 213 × 113 in the tail region of a live zebrafish embryo 96 hpf, assembled from 7 × 7 × 3 independently corrected subvolumes (*c.f.*, movie S4). Scale bar, 30 μm. (**B**) Increasing effectiveness of correction, as seen in orthogonal MIPs from 3 ^m thick slabs extracted from the larger volume, under different scenarios: no AO (left column); AO correction from center subvolume applied globally (center column); and independent AO correction in each subvolume (right) (*c.f*, fig. S11). Insets compare, at higher magnification, the quality of correction at the center subvolume (orange box) vs subvolumes at the periphery of the tail (blue boxes). Subvolume boundaries are shown in white. (**C**) A 5 × 4 × 7 set of measured excitation (left column, top) and detection (left column, bottom) aberrations which, after AO correction, yields diffraction-limited imaging over a 170 × 185 × 135 μm volume (left column, center) in the spine of a zebrafish embryo 30 hpf (c.f., Movie 6). Comparative orthoslices before (center column) and after (right column) AO correction show increased aberration but continued recovery of high resolution at progressively greater depth. Scale bar, 30 μm. (D) Aberration corrected volume renderings over 156 × 220 × 162 μm in the spine of a zebrafish embryo, at three time points from a series of volumes acquired from 30–34 hpf at 30 min intervals (c.f., movie S5), flanked on either side by excitation and detection path aberrations at those time points. To compensate for changing aberrations during development, AO correction followed an interleaved strategy where, after initial correction at all 5 × 5 × 6 subvolumes, additional correction was applied at a different subset of 25 subvolumes (green boxes) at each time point, so that all subvolumes were updated every three hours.

Empirically, the largest aberrations we have seen in zebrafish embryos occur at regions of high curvature between the embryo and the surrounding media, or at regions of rapid refractive index change, such as near the notochord (Fig. 4C, Movie 6). For example, when the LLS penetrates the embryo nearly perpendicular to its surface, the excitation aberration is initially small (red arrow, Fig. 4C, column 1, top). However, after the light sheet passes through the notochord, the light sheet encounters substantial aberration, as seen in both the measured wavefront (green arrow, column 1, top) and uncorrected image (green arrow, column 2, bottom). On the detection side, aberrations increase with increasing depth in the embryo (column 1, bottom and column 2, top to bottom). In addition, substantial aberrations occur when the edge of the embryo is imaged tangentially (yellow arrow, column 1, bottom), so that part of the detection light cone intersects the embryo and part does not.

Provided that the specimen does not shift more than a fraction of the isoplanatic patch size while imaging, a given set of tiled AO corrections can remain valid for hours (Movie 6). However, growth during development can cause an embryo to change its shape, position, or refractive index profile such that new corrections are occasionally needed. Fortunately, these changes often occur on a time scale slow compared to that needed to image even a large FOV by LLSM. In such cases, it is sufficient to update the correction at only a subset of different tiles at each time point, as long as all subsets together encompass all tiles in the FOV before the previous round of corrections becomes inaccurate (Fig. 4D, movie S5). Usually, we choose subsets that broadly cover the FOV, in order to monitor where the aberrations change the fastest.

One region that involves substantial specimen curvature, large spatial variation of refractive index, and gradual aberration change is the eye of the developing zebrafish. We imaged (Fig. 5) a 128 × 150 × 75 μm volume consisting of 4 × 4 × 3 tiles (Fig. 5A, Movie 7) spanning most of the eye of an embryo 24–27 hpf at 6 min intervals in order to study differences in the intracellular organization of various organelles (Fig. 5B, C) at different points in the cell cycle. Transgenic labeling of all plasma membranes allowed us to segment, isolate (Fig. 5D), and characterize each cell by type (Fig. 5E). Skin cells exhibited mitochondria clustered in the perinuclear region, similar to mitochondria seen in flat and thin cultured cells to which they are morphologically similar. Conversely, mitochondria in retinal neuroepithelial (RNE) cells were generally longer, distributed across the length of the cell, and polarized along the same axis as the cell itself. The ER in RNE cells, although broadly distributed, was usually most dense around the nucleus and least dense near the polarized ends. In mitotic cells, however, we again observed (*14*) that the ER remodeled into sheet-like structures concentrated near the PM (Movie 7).

**Fig. 5.**
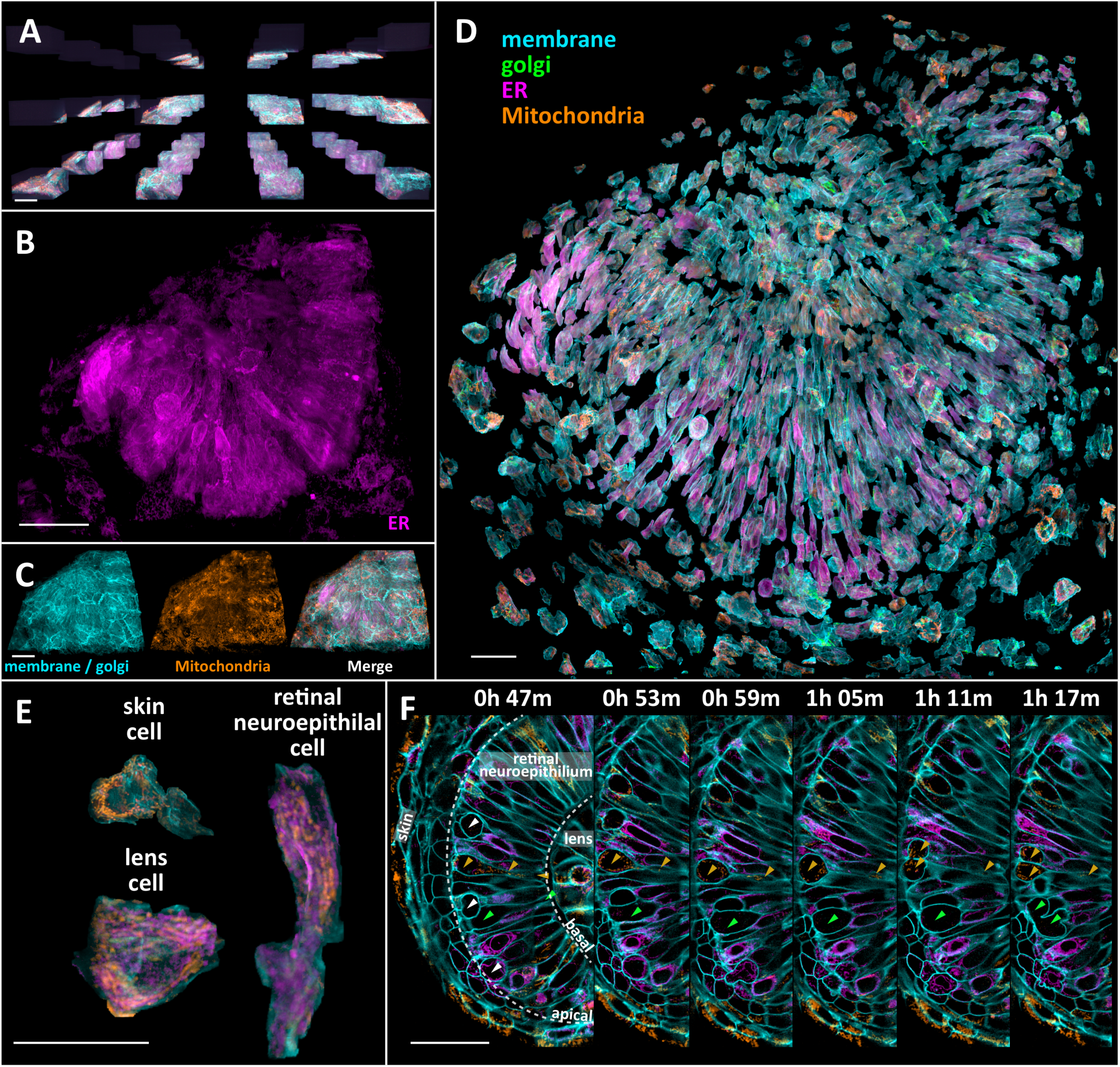
Organelle diversity across the zebrafish eye. (**A**) 4 × 4 × 3 subvolume array used to provide AO correction across the eye of a developing zebrafish embryo at 30 time points from 24.0 to 26.8 hpf (*c.f*, Movie 7). Scale bar, 30 μm. (**B**, **C**) Distribution of three different types of organelles across the 128 × 150 × 75 μm volume assembled from the subvolumes in Fig. 5A. Scale bars, 30 μm. (**D**) Computationally separated cells across the eye, with the three organelles highlighted as shown. Scale bar, 30 μm. (**E**) Cellular and organelle morphologies in cells of three different types within the eye. Scale bar, 30 μm. (F) Orthoslices at six different time points highlighting cell divisions (arrows) at the apical surface of the retinal neuroepithelium and mitochondria (orange arrows) present from the apical to the basal surface in one dividing cell. Scale bar, 30 μm.

Imaging over time (Fig. 5F), we could follow the stages of RNE cell division (green arrows). As reported elsewhere (*15*), we found that, prior to mitosis, the nucleus retracts to the apical side of the retina (leftmost orange arrows, interkinetic nuclear migration), while the cell maintains a thin connection to the basal side (rightmost orange arrows). Despite its narrowness, mitochondria remain in this region. RNE cell divisions then occur at the apical surface (white arrows). It has been found (*16*) that this process is necessary to maintain the integrity of retinal tissue.

## 3D Cell Migration *In Vivo*

*In vivo* 3D migration of a cell in the densely crowded environment of living tissue involves forces, constraints, elasticity and adhesion heterogeneity, and chemical cues not found in the simple 2D environment on a cover glass. Furthermore, cell migration involves intricate and rapid remodeling of membranes, organelles, and the cytoskeleton that requires high spatiotemporal resolution to observe. It is therefore a problem well-suited to the unique capabilities of AO-LLSM.

One such problem involves the wiring of neuronal circuits during development. To help them establish precise connections, axons are tipped with a highly complex and motile structure, the growth cone. This structure functions as both a sensor and a motor, driving the growth of neurites based on environmental cues (*17*). Although its dynamics have been shown to be important for its proper function (*18–20*), its 3D dynamics in an intact animal has been difficult to study because of the lack of imaging techniques capable of resolving the structure with sufficient resolution in all three dimensions *in vivo.*

To address this, we employed AO-LLSM to image growth cones in the spinal cord of a zebrafish embryo in which a subset of newly differentiated neurons expressed stochastic combinations of three different fluorophores via Autobow (*21*) (Fig. 6A, fig. S12) so they could be spectrally distinguished from earlier differentiated neurons expressing only mCherry (e.g., those within the medial neuropil of the reticulospinal tract (magenta arrows, Fig. 6A)). The Autobow-labeled neurons include Rohon-Beard sensory neurons in the dorsal spinal cord (e,g, yellow arrows, Fig. 6A) and interneurons with commissural axons. Imaging a 60 × 224 × 180 μm volume covering more than two spinal segments at 10.4 min intervals from 58 to 70 hpf, (Fig. 6B, Movie 7), we observed that the growth cones of axons migrating in the rostrocaudal direction primarily probed in the direction of their motion (Fig. 6C, top, movie S6), whereas the growth cones of dorsoventrally aligned axons probed across a broader two dimensional fan (Fig. 6D, top). Transverse views (Fig. 6C&D, bottom) revealed that most if not all growth cones of both types were located close to the surface of spinal cord, with their filopodia preferentially aligned parallel to the surface, even though the dorsoventral projecting axons had to pass through the spinal cord to reach its surface. This is consistent with the previous observation that the neurites of late-born V2a ipsilateral projecting interneurons are located laterally to the preexisting ones, forming a layer-like organization based on the age of neurons (*22*), and extends this notion to other classes of spinal neurons. This also raises an interesting possibility that the shape of the growth cone is actively controlled *in vivo* to keep its exploration within a layer of its own age group.

**Fig. 6.**
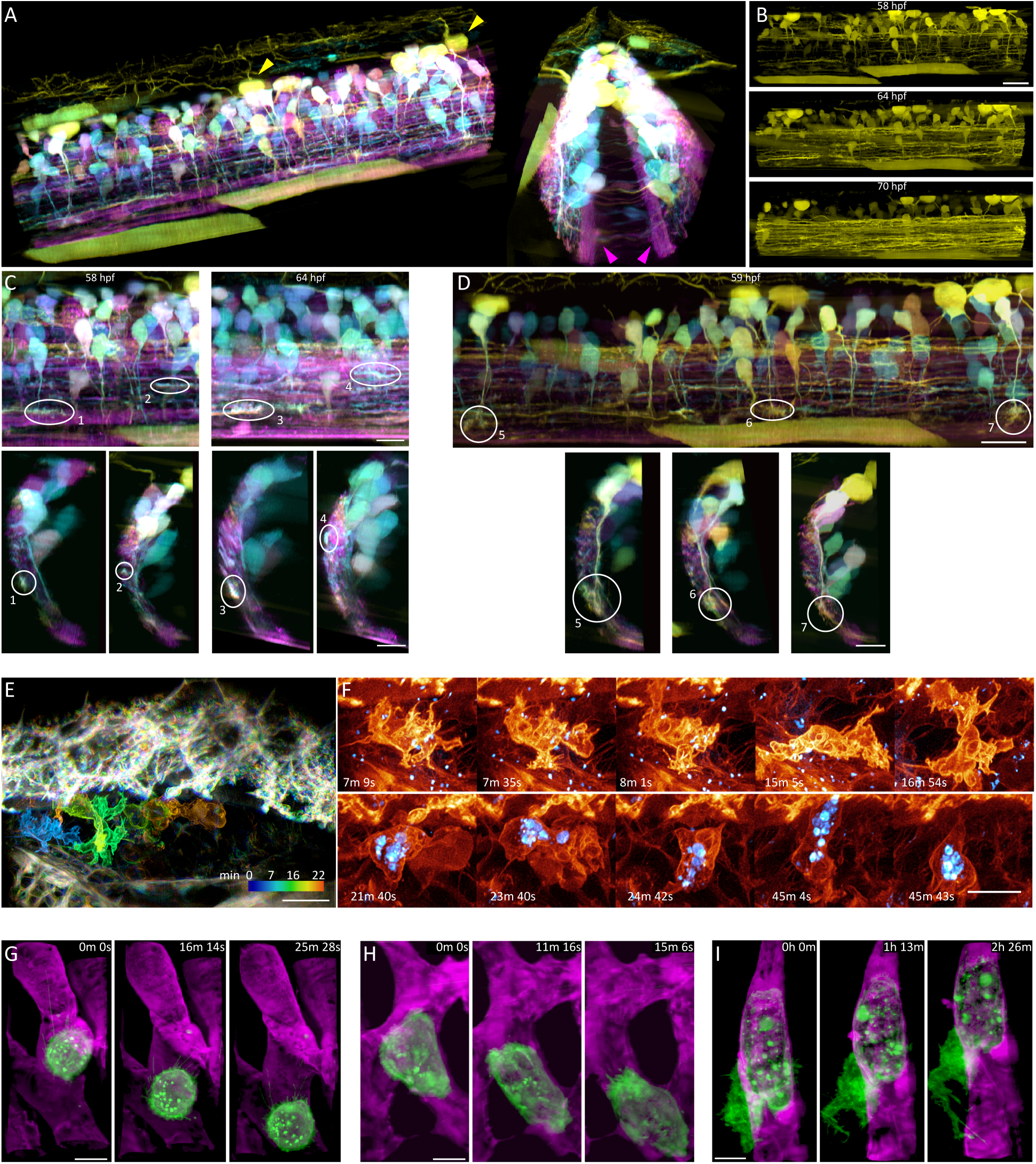
3D cell migration *in vivo.* **(A)** Two views of newly differentiated neurons highlighted by Autobow labeling in a 60 × 224 × 180 section of the spinal cord of a zebrafish embryo 58 hpf. **(B)** Increase in the density of rostrocaudally projecting axons over time. Scale bar, 20 μm. (**C**) Sagittal (top) and transverse (bottom) views of the growth cones of four rostrocaudally projecting axons at 58 or 64 hpf. Scale bars, 10 μm. (**D**) Sagittal (top) and transverse (bottom) views of the growth cones of three dorsoventrally projecting axons at 59 hpf. Scale bars, 10 μm. (**E**) Time-coded color overlay of an immune cell migrating within the perilymphatic space in the inner ear of a live zebrafish embryo 70 hpf, at time points 1–106 from a series of 438 time points at 13 sec intervals (*c.f.*, Movie 9, fig. S12). Scale bar, 10 μm. (**F**) Complex and changing morphologies of two different immune cells (top and bottom rows), one showing internalized dextran particles (blue) (*c.f.*, fig. S13). Scale bar, 5 μm. (**G**) MDA-MB-231 human breast cancer cells (green) rolling in a blood vessel (magenta) within a zebrafish embryo 48 hpf. (**H**) Another MDA-MB-231 cell crawling through a blood vessel. (**I**) A partially extravasated MDA-MB-231 cell, showing an increasingly complex morphology over time (*c.f*, Movie 10, fig. S1). Scale bars, 10 μm in G-I.

Cell migration is also a key aspect of the innate immune system. Neutrophils, for example, must migrate from the vasculature and through the endothelium to reach and engulf infectious targets (*23*). To study this process *in vivo*, we imaged endogenously produced immune cells moving through the perilymphatic space of the ear in a transgenic zebrafish larva expressing the fluorescent protein Citrine in the PM of all cells ~80 hpf (Fig. 6E, Movie 9). Acquiring 438 image volumes at 13 sec intervals allowed us to accurately measure the 3D position and speed (fig. S13A, B) of the cellular center of mass. Across five trials involving different embryos at 22°C, immune cells moved by crawling, adopting a halting search pattern involving frequent changes in direction and speed (fig. S13C-G), from nearly motionless to 8–10 μm/min. In contrast, neutrophilic mammalian HL-60 cells imaged in a collagen matrix at 37°C (*2*) exhibited peak speeds of ~25 μm/min (fig. S13H), perhaps reflective of the lack of local chemoattractive targets and homogeneous ECM, as well as the increased temperature.

Immune cells in the ear were remarkable for their rapidly changing and complex 3D morphologies (Fig. 6F) that would be difficult to observe in detail without the speed and resolution of AO-LLSM. Their surface areas changed as much as 25% in two minutes (fig. S13I) as they remodeled themselves to exhibit a variety of protrusive features, including lamellar sheets, blebs, and short filopodia. Frequently, they also trailed a much longer single filopodium behind them as they migrated to new regions (fig. S14). After injecting fluorescent Texas Red dextran into the heart, we also observed several immune cells containing granules of dextran up to several μm in size (light blue, Fig. 6F and Movie 9, part 1), presumably ingested earlier by phagocytosis. However, these did not noticeably affect the motility of the cell or its ability to navigate through tight interstitial spaces.

Apart from immune cells, 3D AO-LLSM movies (e.g., Movie 9, part 2) of the developing ear region displayed a wealth of cellular morphologies and behaviors, including filopodial oscillations at the dorsal surface of skin cells, gradual inflation of the perilymphatic volume, rapid passage of cells through blood vessels, overlapping lamella at the endolymphatic sac, long and active filopodia on endothelial cells, and cellular rearrangements in the hindbrain. With adaptive optics, the spatiotemporal resolution and non-invasiveness we achieved was comparable to that we obtained when imaging cultured cells in our original LLSM (*2*), allowing us, for example, to follow the detailed morphological changes in a single endothelial cell lining the hindbrain over the entire course of its division (fig. S15 and Movie 9, part 2).

As a final example of 3D migration, during cancer metastasis circulating tumor cells (CTCs) extravasate and seed new tumor formation at sites distant from the primary tumor (*24*). Extravasation has been studied extensively *in vitro*, but little is known about the process in *in vivo*, due to the highly dynamic nature of cells in circulation, and the low density of CTCs in the vasculature. A long-standing hypothesis based primarily from *in vitro* studies is that CTCs co-opt mechanisms used by leukocytes to extravasate at sites of inflammation (*24*). During this process, leukocytes progress through an adhesion cascade and extravasate in a three-step process (*25*). Circulating leukocytes initially weakly adhere to the endothelium through selectin-mediated adhesion, which causes them to roll slowly along the blood vessel in the direction of blood flow. Eventually the leukocytes stop rolling and switch to an integrin-dependent crawling behavior along the endothelial wall, ultimately penetrating the endothelium during transendothelial migration. Determining whether CTCs follow this behavioral pattern *in vivo* requires rapid high resolution imaging.

To visualize CTC extravasation *in vivo*, we used a zebrafish xenograft model (*26*). Human breast cancer cells (MDA-MB-231) with a membrane label were injected into the vasculature of two-day old zebrafish embryos transgenic for an endothelial reporter (*kdrl:gfp*). We observed all three leukocyte behaviors in the cancer cells. First, we recorded MDA-MB-231 cells rolling through the blood vessels (Fig. 6G). Leukocyte rolling occurs through the formation of microvilli, which form weak adhesive catch bonds to the endothelium through selectin mediated adhesion. In MDA-MB-231 cells, we also observed such microvilli adhering to the endothelium, stretching several microns as the cell moves with the flow of blood before recoiling as the weak adhesion is broken (Fig. 6G and Movie 10, part 1). Second, we visualized MDA-MB-231 cells crawling along the endothelium (Fig. 6H & Movie 10, part 2). And lastly, we observed a MDA-MB-231 cell actively engaged in transendothelial migration, with the portion of the cell outside the blood vessel projecting actin-rich extensions into the surrounding tissue (Fig. 6I and Movie 10, part 3) as the area of the cell increased by ~50% over two hours (fig. S16). Together, these movies provide a high-resolution view of *in vivo* cancer cell extravasation that indeed recapitulates the three steps of leukocyte adhesion, crawling, and transendothelial migration.

## Discussion

By combining lattice light sheet microscopy with two independent channels of adaptive optical correction, AO-LLSM enables minimally invasive high speed 3D imaging of subcellular dynamics within optically challenging living specimens while still maintaining diffraction-limited resolution, even over large fields of view. Furthermore, it corrects not only sample induced aberrations, but also those introduced by mounting / immersion media (*e.g.*, Fig. 1C) or imperfections in the optical path through the microscope. It therefore can provide practical 3D resolution exceeding that of nominally higher numerical aperture (NA) confocal or spinning disk microscopes, even in the comparatively benign optical environment encountered when imaging isolated adherent cells on cover slips.

This performance does not come without caveats, however. Because the fluorescence induced by the light sheet is captured with widefield optics, only weakly scattering specimens can be imaged. In addition, extremely sparse and/or weakly emitting fluorescent targets may require co-labeling with a second, brighter color channel to provide sufficient guide star signal (~100 photons or more per cell of the Shack-Hartmann sensor) for accurate wavefront measurement. Highly absorbing structures such as large blood vessels or melanin bodies within the detection light cone can block guide star light from reaching enough cells of the sensor for accurate wavefront measurement, although it may be possible to develop wavefront reconstruction algorithms more robust when faced with such missing information. Even when measured accurately, wavefront aberration can vary considerably across the specimen, and this variation currently can only be determined empirically for each specimen type and developmental stage to determine how to subdivide the desired image volume into tiled isoplanatic subvolumes of relatively uniform aberration. Fortunately, such tile maps tend to be consistent between specimens of the same type and age, given similar mounting geometries. Finally, specimens imaged after muscle development must be anesthetized and immobilized, or else a new correction must be measured and applied whenever sample motion exceeds the size of a given isoplanatic patch.

Notably, all but one of the above examples involved imaging subcellular dynamics within zebrafish embryos. While we have shown that we can achieve substantial gains in imaging performance in both *Caenorhabditis elegans* larvae (fig. S17, movie S7) and *Arabidopsis thaliana* leaves (fig. S18, movie S8), *Danio rerio* represents an ideal model system in which to study cell and developmental biology *in vivo*, as it is a rapidly developing transparent vertebrate that is amenable to genetic manipulation. Furthermore, zebrafish embryos are small enough that most regions are optically accessible far into development, yet large enough to exhibit smoothly varying refractive index profiles that result in isoplanatic patch sizes comparable to imaging fields typical of LLSM. In contrast, *C. elegans* larvae and adults exhibit larger and more rapid spatial variations in refractive index, particularly near the gut, that can require a denser mesh of AO corrections, despite its reputation as an optically tractable model organism.

Of course, conventional light sheet microscopy using weakly focused Gaussian beams is also susceptible to aberrations, and would therefore also benefit from AO correction (*4, 5*). However, conventional systems typically cover much wider fields of view and often operate at greater depth in larger organisms, such as in applications involving functional imaging of whole neural circuits (*27*) or *in toto* cellular tracking during development (*28*). They therefore usually image over regions much larger than a single isoplanatic patch, making it difficult to retain even only cellular-level resolution at all locations, and compromising the accuracy and resolution of approaches based on multi-view fusion (*29–31*). A single AO correction would provide at best only partial correction, and a tiled AO approach, such as we use with LLSM, would negate the high speed, large field advantages of the conventional light sheet microscopy. On the other hand, simultaneous full field AO correction would likely require multi-conjugate adaptive optics (*32*) using several guide stars and wavefront modulation elements operating in concert, substantially increasing cost and complexity.

Perhaps the greatest challenge of AO-LLSM involves mining the immense and complex data it produces to extract as much biological insight as possible. Figs. 6A-D, for example, represent a minute fraction of a 0.62 TB raw data set which first had to be deconvolved, creating a second copy, and then imported into 3D visualization software, generating a third. We deconvolve and store data in real time, but importation and visualization can take many hours, preventing meaningful real-time feedback on whether the desired biological structure and dynamic process of interest is being optimally recorded. If history is any guide, problems of petabyte scale data storage and visualization at reasonable cost will yield to continued advancements in commercial hardware, but problems of image analysis and meaningful quantification of data at this scale may prove far less tractable. While we have demonstrated quantification on a smaller scale through single particle (Fig. 2D, E, fig. S8) and single cell (fig. S13) tracking, segmentation (Figs. 2B, 3A, & 5D), and measurement of area and volume (Fig 3E, F, figs. S9, S16), the diversity of questions that can be asked when modern genetic and pharmacological tools are combined with high resolution 5D *in vivo* data spanning hundreds cells over many hours will demand bioinformatics expertise, machine learning, and custom algorithm development on an unprecedented level. Nevertheless, such efforts promise to offer insights into how cells harness their intrinsic variability to adapt to different physiological environments and to reveal the phenotypic diversity of organelle morphologies, intracellular dynamics, extracellular communication, and collective cell behavior across different cell types, organisms, and developmental stages.

## Materials and Methods

### Lattice Light Sheet Subsystem

The lattice light sheet excitation path of the AO-LLSM was designed as described previously (*2*). Noted here are the changes introduced in the AO-LLSM. The collinear laser beams from the combiner were first expanded using a pair of cylindrical lenses and aligned such that up to three different wavelengths illuminated three vertically separated thin stripes on spatial light modulator SLM, (Holoeye, PLUTO-Vis-014 1920 × 1080 pixels, fig.S1). As a greyscale phase modulation device, SLM was introduced to not only create the light sheet but to correct sample-induced aberrations as well. The diffraction orders reflected from SLM were then filtered using annular mask MSK, (Photo Science Inc) as before and conjugated to galvanometer scanning mirrors G3 and G4 (3 mm mirror, Cambridge Technology, 8315H) to scan the light sheet along the x and the z axis. During imaging, different offset voltages were applied to the z galvo to sequentially realign the light sheet from each laser to the same plane within the specimen. Sample plane conjugate resonant galvanometer RG (Electro-Optical Products Corp. 7 × 8 mm, SC-30) was also added prior to the excitation objective (Special Optics, 0.65 NA, 3.74 mm WD) to wobble the light sheet in the xy plane and thereby minimize stripe artifacts due to localized absorbing or scattering objects in the specimen. The fluorescence generated in the excitation plane was collected with detection objective DO (Nikon, CFI Apo LWD 25XW, 1.1 NA, 2 mm WD) and reflected off deformable mirror DM (ALPAO 97–15) conjugate to the rear focal plane of DO before being imaged at sCMOS camera CAM 1 (Hamamatsu Orca Flash 4.0 v2). Complete details of the optical design are given in supplementary note 1.

### Adaptive Optics Subsystems

In principle, independent AO corrective systems are needed for excitation and detection in LLSM, since light traverses different regions of the specimen in each case and hence is subject to different aberrations. However, given that: a) aberrations decrease quickly with decreasing numerical aperture (NA) (*3*); b) we use at most 0.6 NA for excitation, versus 1.1 NA for detection in LLSM; and c) only aberrations within a narrow annulus at the rear pupil of the excitation objective will affect a lattice light sheet, it is not obvious that AO correction of the light sheet itself is necessary. To check, we simulated the effect of aberrations consisting of random combinations of the 55 lowest order Zernike modes up to a root mean square (RMS) amplitude of two wavelengths (l). We found (fig. S19) that aberrations at this level could expand a 0.7 μm thick lattice light sheet to as much as 20 μm, and displace it perpendicular to its plane by up to ±8 μm, indicating that correction of excitation as well as detection is essential.

Hence, during aberration measurement, light from Ti:Sapphire ultrafast pulsed laser 2PL (Coherent Cameleon Ultra II) was ported to either the excitation or detection arm by switching galvanometer SG1 (fig. S1). In either case, TPEF generated within the specimen by scanning the guide star focused by EO or DO was collected by the same objective, descanned (*7*) and sent to homebuilt Shack-Hartmann wavefront sensor ESH or DSH, each consisting of a square microlens array (Edmund Optics) focused onto an EMCCD camera (Andor iXon). Corrective wavefronts were then applied to SLM or DM as described in supplementary notes 3 or 2 respectively. Further hardware details are given in supplementary note 1.

Autofocus measurement was achieved by viewing, side-on through EO, both the light sheet fluorescence and the plane of fluorescence generated by guide star TPE excitation through DO on camera CAM4, and correcting for any displacement between them as outlined in supplementary note 5.

### Control Electronics

Control electronics were similar to those described previously (*2*), with the primary changes concerning the SLM and computer. In the new configuration, synchronization is simplified because the SLM now displays fixed images and only updates when the excitation correction is changed via software command between volume scans. A Field-Programmable Gate Array card (FPGA, National Instruments, PCIe-7852R) serves as the master clock supplying camera triggers and control voltage waveforms during image acquisition. Additional PCIe slots are needed to accommodate the added cameras and DM, and thus the control computer has been upgraded to a SuperMicro 4027GR-TRT with dual Intel E5–2670 processors.

### Zebrafish immobilization, mounting and imaging conditions

Zebrafish embryos were paralyzed with ~1 ng of α-bungarotoxin protein injected prior to imaging (*33*) or anesthetized using tricaine (0.16mg/ml) for 15 minutes. 12 mm diameter glass coverslips were pre-cleaned as follows: ~ 20 coverslips were placed in a 50 ml Falcon tube containing 0.1M NaOH and the tube placed in a sonicator for 15 min, followed by at least 5 consecutives washes with Mili-Q water and then immobilized in the sample holder using superglue. An agarose holder containing narrow groves for mounting the embryos was created by solidifying a few drops of 0.5–2% (wt/wt) low melting agarose between the coverslip and a mold containing ridges. For the immune cell experiments, larvae were mounted in custom volcano mounts made with a 3-D printed mold (https://www.shapeways.com/shops/megason-lab). A homemade hair-loop was used to position the embryo in the mold, which was then stabilized with a thin layer of agarose made by applying on top of the immobilized embryo ~10-20^l 1% low melting agarose at 37–40°C and then wicking the excess. After solidification, the sample holder was bolted onto a three-axis set of sample stages (Attocube, ECS3030 for x and y, ECS3050 for z) and submerged in a sample bath containing ~8 ml of 1x Danieau buffer. This assembly was then raised by a motorized actuator (Newport, LTA-HS Actuator, Integrated with CONEX-CC Controller, CONEX-LTA-HS) until EO and DO were immersed in the media. The sample stages then positioned the desired field of view to the mutual focal point of the objectives. Detailed imaging conditions for each experiment discussed in the paper, including excitation power, imaging time, image, tile and voxel sizes, fluorophores and proteins, etc., are in table S1. Additional preparation conditions are discussed in supplementary note 6.

### Image Processing and Visualization

All data acquired using the AO-LLSM were corrected for intensity variation across the light sheet, deconvolved using an experimentally measured PSF for each emission wavelength, and corrected for photobleaching as noted in table S1 and as described (*2*). Multi-tile subvolumes were stitched as indicated (table S1) using either the Grid/Collection Stitching plugin in Fiji (*34*), or a Gradient-Domain stitching routine (*35, 36*) to merge and smooth the boundaries between adjacent tiles by matching the low spatial frequency components. All processed data sets were visualized using Amira (Thermo Fisher Scientific), Imaris x64 8.4 (Oxford Instruments), or Vision4D (Arivis) for 5D volumetric rendering.

The cell boundaries within different tissues of zebrafish embryos expressing plasma membrane markers were automatically segmented using Automated Cell Morphology Extractor (ACME) (*37*). Using the segmented masks, a novel computer-aided displacement (CAD) approach was implemented to facilitate the visualization and inspection of the morphology and dynamics of individual cells contained within a portion of the living tissue of the zebrafish embryo. Each cell within a given 3D volume was computationally separated from the others using the following steps: (*1*) determine all cell boundaries as described above to detect multicellular boundaries in zebrafish embryos; (*2*) generate a 3D ROI for each cell using a dilated segmentation mask (sigma = 2); (*3*) displace the centroid position of the cell by multiplying the distance of the cell centroid to the center of the imaged 3D volume by a constant expansion factor. Thus, cells farther from the center of the 3D volume are displaced more than the cells closer to the volume center; (*4*) sequentially repeat step (*3*) with incremental expansion factors.

### Wavefront Reconstruction for Visualization

Visualized wavefronts (e.g., Fig. 4C, D, Movies 6–8) were calculated from valid Shack-Hartmann spots using a zonal wavefront reconstruction (*38*) in MATLAB. Invalid points in the Shack-Hartmann image are recovered by smoothly interpolating using the neighboring points via the gridfit function in MATLAB. The waveforms were then least-squares fitted with the first 55 Zernike modes. The tip, tilt, and defocus modes were set zero. Since the wavefront fit is unconstrained at the edges, the wavefront is mostly clearly plotted when just the inner 80% of the rear pupil diameter is shown. The wavefronts were rendered using Amira 6.3 at the tiled locations where the measurements and corrections took place.

## Supplementary Materials

Supplementary Notes

Figures S1-S20

Table S1

Movies S1 to S8

Full Reference List

## Acknowledgments

We thank the Shared Resource teams at the Janelia Research Campus for their skill and dedication in specimen handling and preparation, and the Instrument Design and Fabrication team for their manufacturing expertise. We also gratefully acknowledge the support of the Janelia Visitor Program. TLL, DEM, VS, JS, MK, EMM, and EB are funded by the Howard Hughes Medical Institute (HHMI). TK and SU are funded by grants from Biogen, Ionis Pharmaceuticals and NIH R01GM075252 to T.K. SU is a Fellow at the Image and Data Analysis core at Harvard Medical School and thanks H. Elliott, D. Richmond and R. Gao for discussions and the MATLAB code repository received from the Computational Image Analysis Workshop supported by NIH grant GM103792. SS is funded by a LSI start-up grant, University of Exeter. KRM, IAS, ZMC, TWH, and SGM were supported by R01DC015478. DQM is funded by the NIH (5R00CA154870–05, 1R01GM121597–01). DQM and BLM are funded by the Carol M. Baldwin Foundation and are Damon Runyon-Rachleff Innovators supported (in part) by the Damon Runyon Cancer Research Foundation (DRR-47–17). BLM is also funded by the NSF (IOS1452928). D.H. is a Pew-Stewart Scholar for Cancer Research supported by the Pew Charitable Trusts and NIH R01-CA196884. D.D. was supported by a Human Frontier Science Program Fellowship and D.G.D. was supported by National Institutes of Health Grant R35GM118149. Portions of the technology described herein are covered by U.S. Patent 7,894,136 issued to EB and assigned to Lattice Light, LLC of Ashburn, VA, U.S. Patents 8,711,211 and 9,477,074 issued to EB and assigned to HHMI, U.S. Patent application 13/844,405 filed by EB and KW and assigned to HHMI, and U.S. Patent 9,500,846 issued to EB and KW and assigned to HHMI.

## Movie Captions

**Movie 1.**
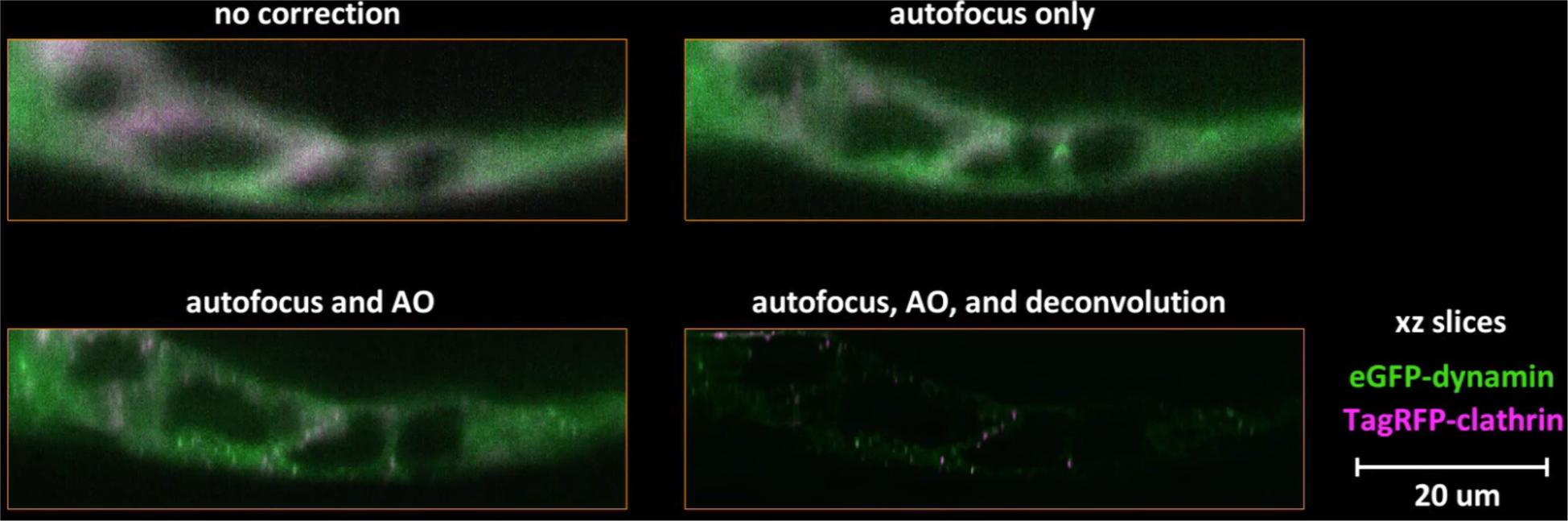
Endocytosis in a human stem cell derived organoid. Gene-edited clathrin (magenta) and dynamin (green) before and after adaptive optical correction and deconvolution, showing comparative xy and xz orthoslices, comparative volume renderings, and post-correction tracking of the motion and lifetimes of individual clathrin coated pits and coated vesicles over 120 time points at 1.86 sec intervals (*c.f.*, Fig. 1C, fig. S5).

**Movie 2.**
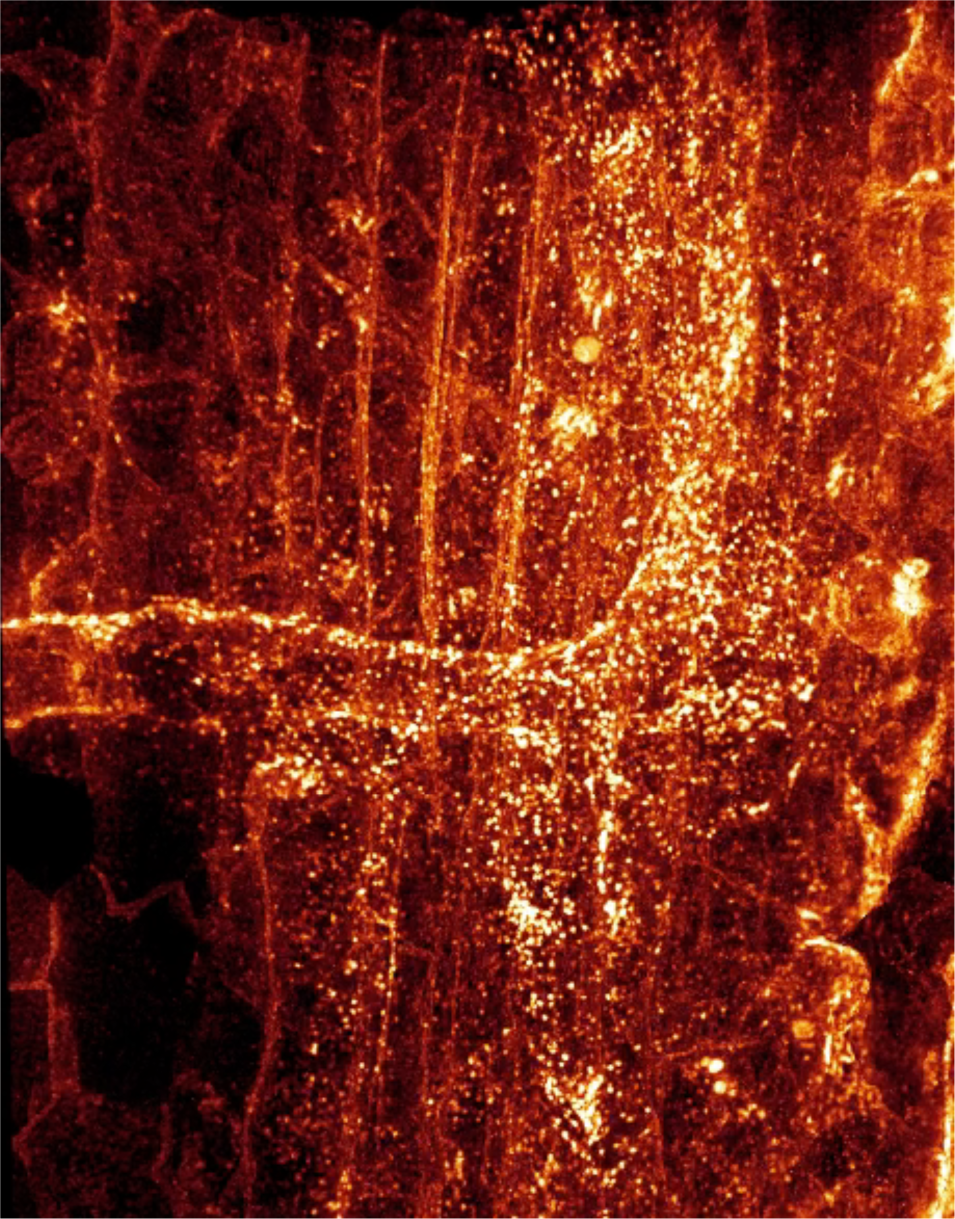
Clathrin-mediated endocytosis *in vivo.* Dynamics of CCPs and CCVs over 15 min at 10 sec intervals in a 75 × 99 × 40 μm volume from the dorsal tail region of a zebrafish embryo 80 hpf. Segmented cells reveal brighter clathrin puncta at the vascular endothelium than at muscle fibers (*c.f.*, Fig. 2A).

**Movie 3.**
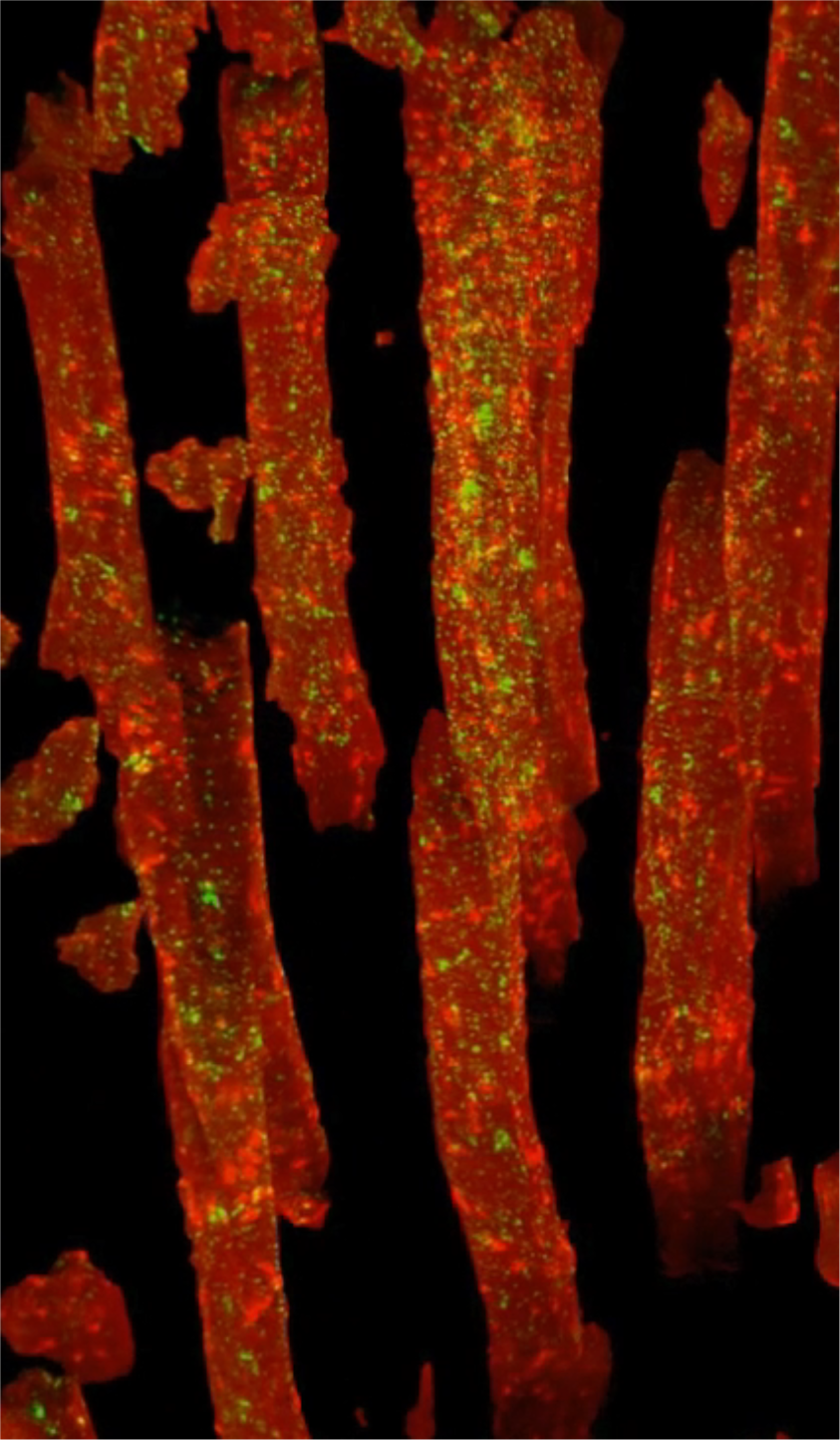
Clathrin localization in muscle fibers. Plasma membranes (red) and clathrin (green) in the tail of a zebrafish embryo 5G-55 hpf, showing: xy and xz orthoslices before and after AO correction and deconvolution; dynamics of individual CCPs and CCVs at and between t-tubules; large clathrin clusters and small clathrin puncta in volume rendered and segmented cells; and tracked CCPs and CCVs in a segmented cell (*c.f.*, Fig. 2B-E, fig. SS, movie S3).

**Movie 4.**
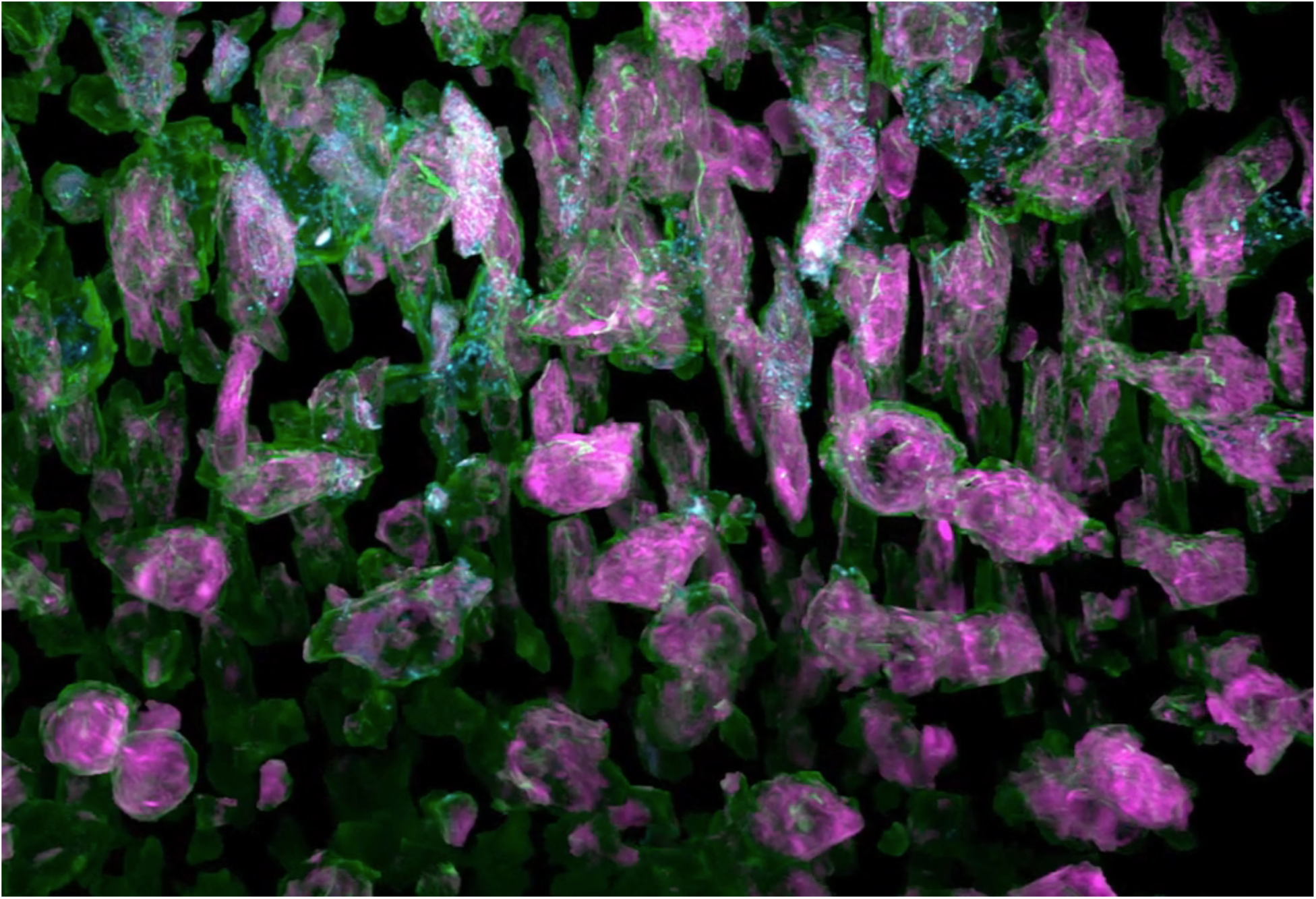
Subcellular imaging of organelle dynamics in the early zebrafish brain. Dynamics of plasma membranes and trans-Golgi (green), endoplasmic reticulum (magenta or red), and mitochondria (cyan) within neural progenitor cells over 200 time points at 44 sec intervals from 14.0 hpf, showing: complexity within the original 70 × 35 × 35 μm image volume; cross-sectional slab views through cells; sequential division of adjacent cells; segmentation and separation of all cells; and morphological changes to organelles during mitosis in one such cell (*c.f.*, Fig. 3A, B, figs.S9, S10).

**Movie 5.**
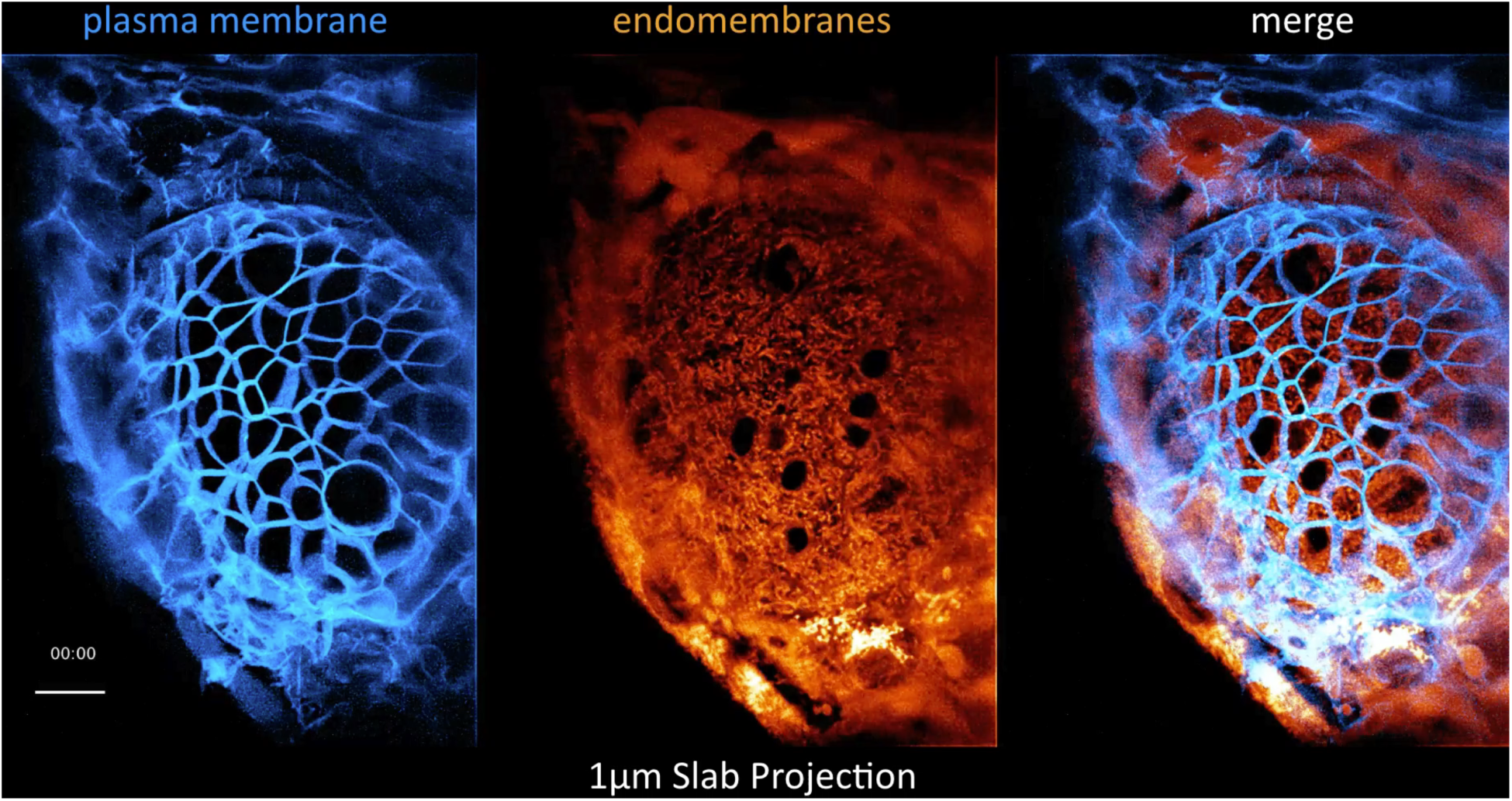
Membrane dynamics in the zebrafish eye. Plasma membranes (blue) and the endomembrane system (orange) 30 hpf viewed as *xy* orthoslices, cell divisions in a 1 μm thick slab, and volume rendered PM dynamics across the eye at 43.8 sec intervals for 200 time points (*c.f.*, Fig. 4C-E).

**Movie 6.**
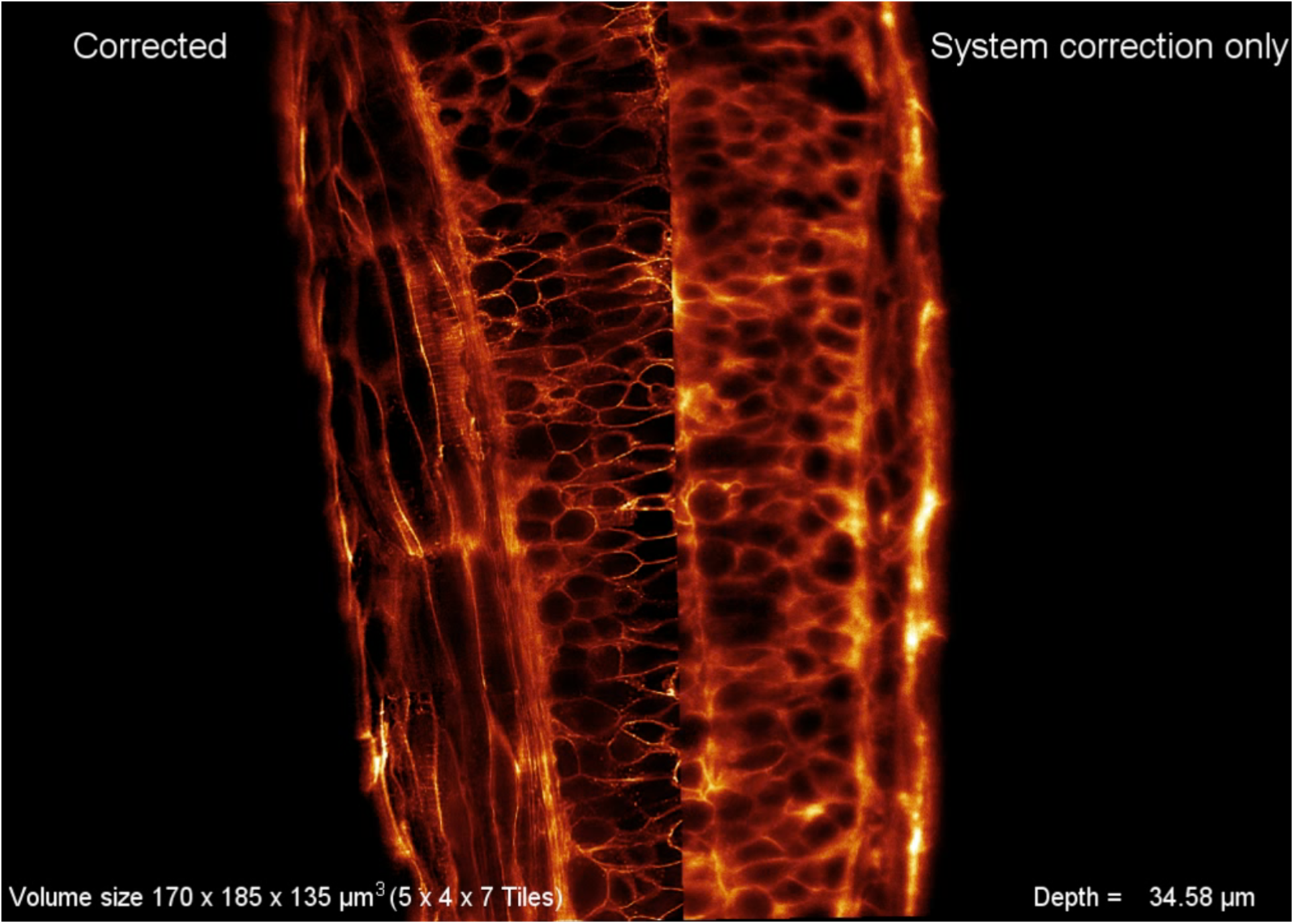
Tiled AO correction for imaging large volumes. 170 × 185 × 135 μm volume from the dorsal surface to the notochord in a pan-membrane labeled zebrafish embryo showing: increasing aberration but continued full correction at increasing depth; corrective excitation and detection wavefronts in each of the 5 × 4 × 7 array of tiled isoplanatic subvolumes required to achieve full correction over the complete volume; and four views of membrane dynamics within the complete volume from 30 to 39.5 hpf, imaged at 7.5 min intervals (*c.f.*, Fig. 4C).

**Movie 7.**
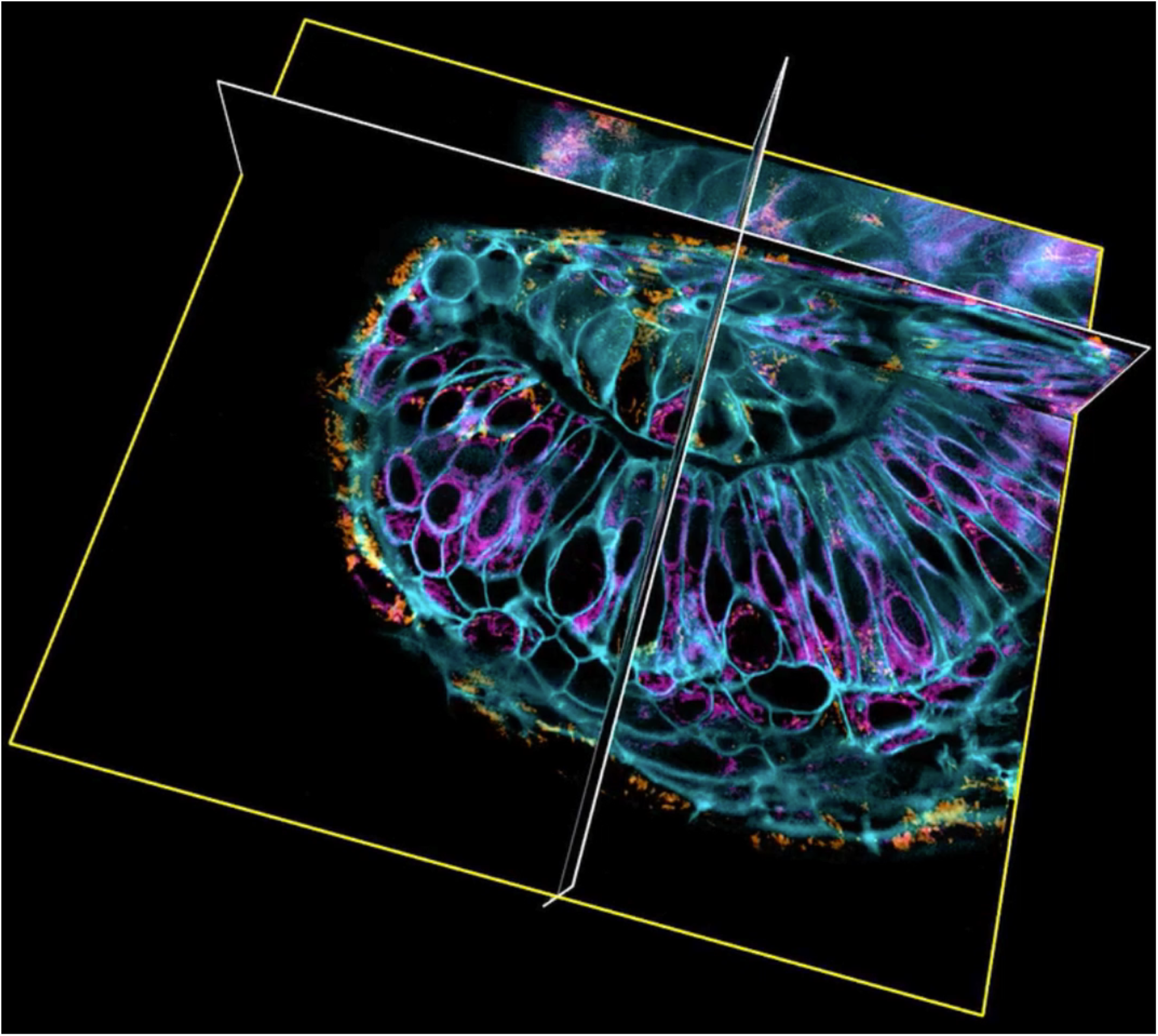
Organelle dynamics across the zebrafish eye. Plasma membranes (cyan), trans-Golgi (green), ER (magenta) and mitochondria (brown) across a 128 × 150 × 75 volume assembled from 4 × 4 × 3 subvolumes showing: orthoslices in a single subvolume; volume rendered subvolumes before assembly into the combined volume; organelles in the combined volume; dynamics over 30 time points from 24.0 to 26.8 hpf in a 1 μm thick slab through the combined volume; dynamics in perpendicular orthoslices; organelle morphologies in different cell types in the segmented and expanded image volume (*c.f.*, Fig. 5).

**Movie 8.**
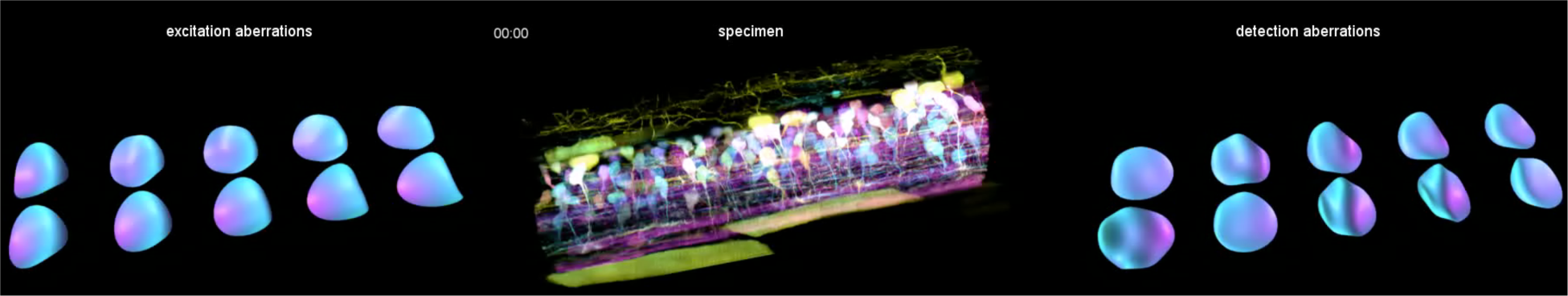
*In vivo* imaging of spinal cord neural circuit development. Autobow labeled, newly differentiated neurons expressing stochastic combinations of three fluorophores in a zebrafish embryo showing: corrective excitation and detection wavefronts in 5 × 2 subvolumes, with scrolling updates at one subvolume (green box) per time point; AO corrected orthoslices and volume rendered views in each color channel 58 hpf; and axon pathfinding in each color channel from 58 to 70 hpf (*c.f*, Fig 6A-D, fig. S12, movie S6).

**Movie 9.**
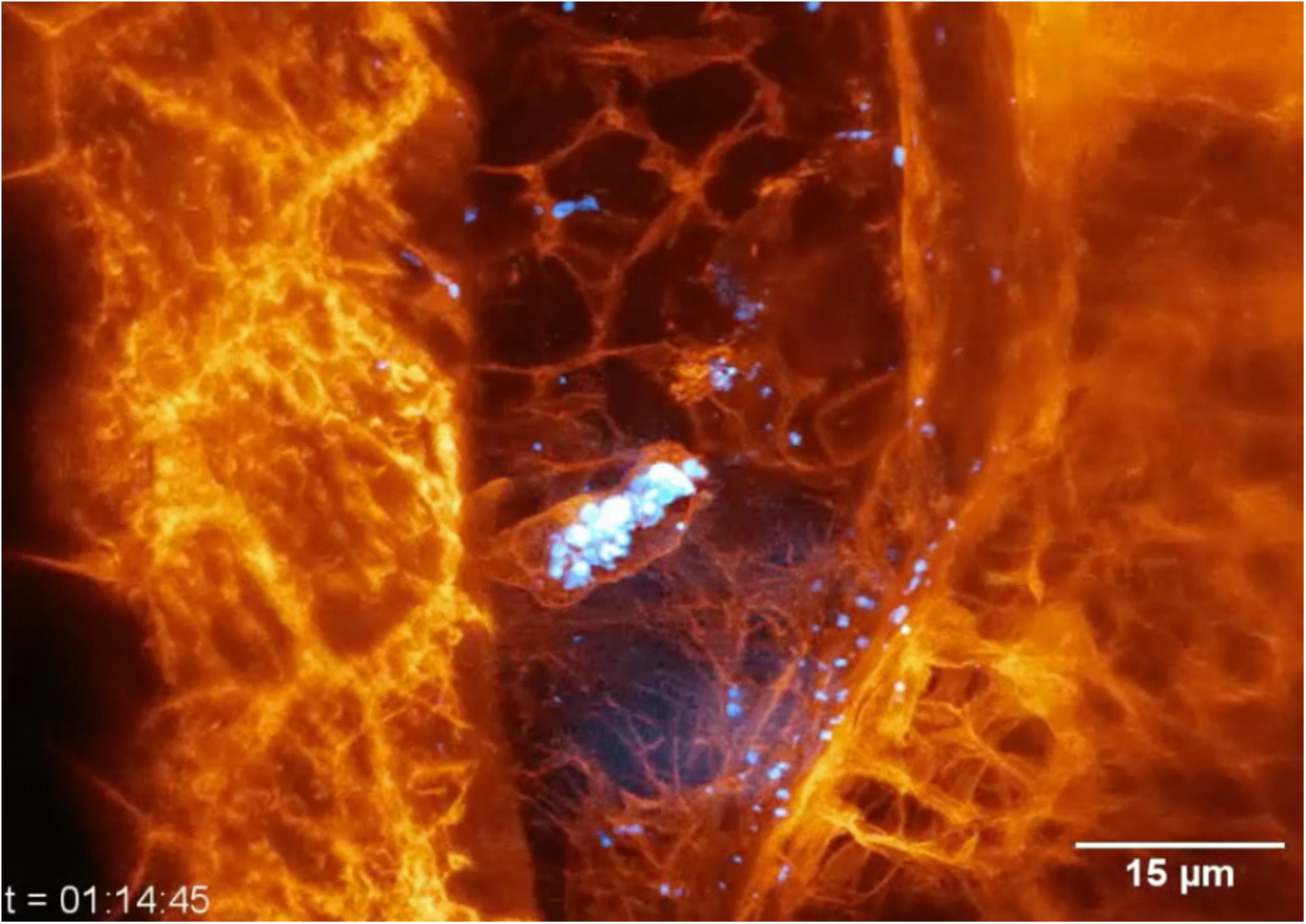
Immune cell migration in the zebrafish inner ear. Immune cells within the perilymphatic space of the inner ear of several zebrafish embryos 80 hpf showing: MIP view of two immune cells (orange), one of which has ingested dextran particles (blue), before and after AO plus deconvolution for 438 time points at 13 sec intervals; volume rendered view in another embryo, showing a migrating immune cell and a dividing endothelial cell; and tracking of the position and velocity of an immune cell in a third embryo (*c.f.*, Fig. 6E,F, figs. S13–15).

**Movie 10.**
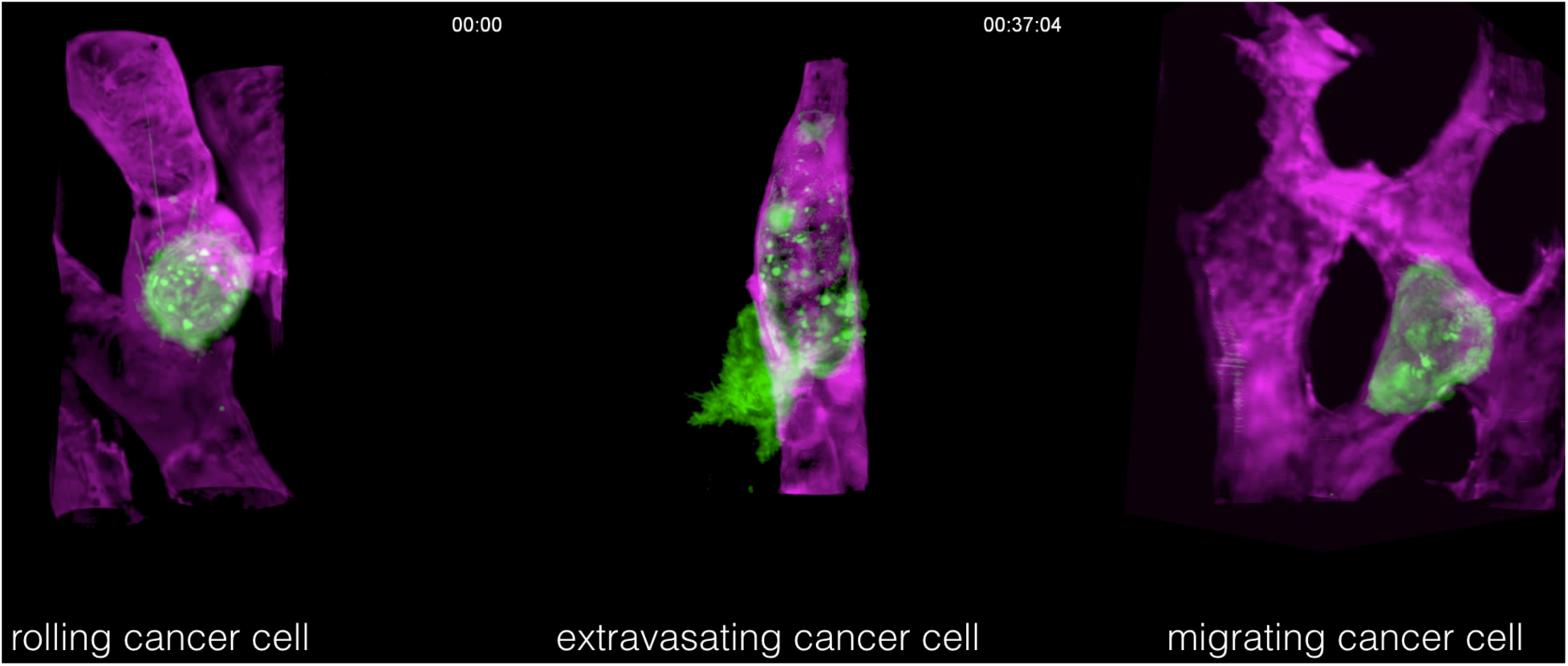
Cancer cell migration in a zebrafish xenograft model. MDA-MB-231 human breast cancer cells (green) exhibiting three different forms of motion within the vasculature (magenta) of different zebrafish embryos: rolling within a blood vessel while extending long, adhesive microvilli; crawling while conforming to the shape of a blood vessel; and partial extravasation from a blood vessel (*c.f.*, Fig. 6G-I, fig. S16).

